# A mixer tap mechanism in *Mycobacterium tuberculosis* regulates a critical metabolic node in response to nutrient availability

**DOI:** 10.64898/2026.04.29.721766

**Authors:** Evelyn Yu-Wen Huang, Katherine J. Jeppe, Jingsong Zhou, Christopher K. Barlow, Jemila C. Kester, Anna K. Coussens, Ivanhoe K.H. Leung

## Abstract

*Mycobacterium tuberculosis* persists within the host by exploiting metabolic flexibility underpinned by the tricarboxylic acid (TCA) cycle-glyoxylate shunt junction, where isocitrate dehydrogenase (ICD) and isocitrate lyase (ICL) compete to direct carbon flux between energy production and carbon conservation. Yet, how nutrient availability regulates flux partitioning at this junction is unknown. We show that among the four gatekeeping isoforms of isocitrate lyase (ICL1/2) and isocitrate dehydrogenase (ICD1/2), only *icl1* transcription substantially responds to carbon substrate switching. Systematic metabolite screening revealed that pyruvate, oxaloacetate, and glyoxylate are novel allosteric activators of ICD2, while concurrently inhibiting ICL1/2 at their physiological concentrations. To resolve how these regulatory layers integrate, we developed an experimental ^1^H NMR-based flux assay that combines recombinant enzymes with metabolomics-informed metabolite conditions to quantify flux partitioning under defined states. This systems-level framework reveals that flux partitioning emerges from coordinated balancing of enzyme abundance and protein-metabolite effector combinations that shift in a time-dependent, nutrient-responsive manner. This dynamic, multi-layered regulatory framework, analogous to a ‘mixer-tap’ mechanism, offers new strategies for therapeutic disruption of *M. tuberculosis* metabolism.

## Introduction

Asymptomatic persistent infection within immunocompetent individuals is a hallmark of *Mycobacterium tuberculosis*, the primary causative agent of tuberculosis^1–3^. Underpinning this persistence is the bacterium’s remarkable metabolic versatility^4–6^. In contrast to the diauxic growth favoured by most bacteria^7,8^, *M. tuberculosis* has evolved a co-metabolic strategy that enables the simultaneous utilisation of glycolytic (e.g. glucose) and non-glycolytic (e.g. fatty acids and cholesterol) carbon sources^9,10^. This supports intracellular survival and virulence in environments with depleted and fluctuating nutrient availability^11,12^. However, the molecular mechanisms that coordinate this metabolic behaviour are poorly understood. This limits efforts to develop new therapies targeting essential metabolic pathways^13,14^.

While both glycolytic and non-glycolytic substrates can support energy generation, growth on non-glycolytic carbon sources requires carbon conservation to replenish intracellular glucose pools via gluconeogenesis^15^. This trade-off between energy generation and carbon conservation to support biosynthetic demands is managed at the junction between the tricarboxylic acid (TCA) cycle and the glyoxylate shunt (**Fig. 1a**)^16,17^. The TCA cycle functions as a metabolic hub, generating energy and electron carriers from carbon substrates, while the glyoxylate shunt conserves carbon through the bypassing of two decarboxylation steps in the TCA cycle, redirecting flux toward gluconeogenesis and biomass production^16,17^. Consistent with this central role, the glyoxylate shunt is essential for growth and virulence on non-glycolytic carbon sources^2,18^, and perturbation of flux partitioning between these pathways inhibits growth of the related species *M. bovis* BCG^19^.

**Figure 1.**
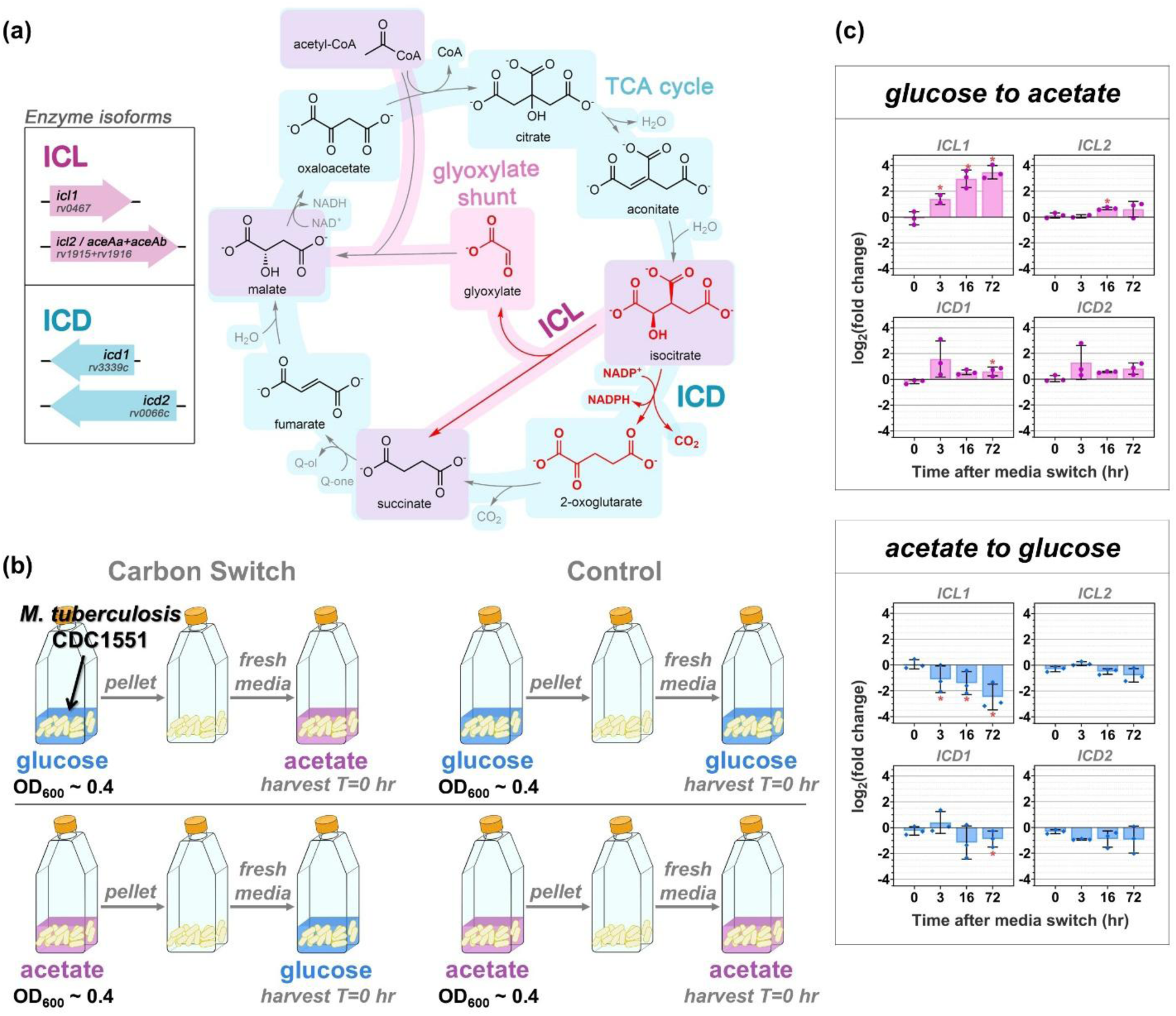
Carbon substrate switching selectively regulates *M. tuberculosis* ICD and ICL isoform gene expression. **(a)** The tricarboxylic acid (TCA) cycle and the glyoxylate shunt. The partitioning of isocitrate between the two pathways is highlighted in red. The reactions catalysed by isocitrate dehydrogenase (ICD) and isocitrate lyase (ICL) are indicated in bold. The ICD and ICL isoforms and their respective gene Rv accession numbers are shown on the left. The TCA cycle (in blue) oxidises acetyl-CoA to generate reducing equivalents that can be converted to high-energy intermediates through oxidative phosphorylation. The glyoxylate shunt (in pink) bypasses the two decarboxylation steps in the TCA cycle. In *M. tuberculosis*, the conversion of 2-oxoglutarate to succinate can occur through different pathways^55–57^, so it is illustrated in this figure as a single reaction without the succinyl-CoA intermediate that is common in many other organisms. Q-one and Q-ol stands for menaquinone and menaquinol, respectively. **(b)** Design of the carbon substrate switching experiment for gene expression analysis in *M. tuberculosis* CDC1551. **(c)** Gene expression changes are calculated relative to the initial time point immediately after the media switch (0 hr). Fold changes were normalised to matched control cultures switched to fresh media maintaining the same carbon substrate. Error bars represent s.d. from three biological replicates. Asterisks (*) indicate statistically significant differences (p < 0.05) relative to each matched time-point control using the Student’s t-test corrected for multiple comparisons using the Holm-Sidak method.

At this junction are two gatekeeping enzymes: isocitrate dehydrogenase (ICD) in the TCA cycle^20^, and isocitrate lyase (ICL) in the glyoxylate shunt (**Fig. 1a**)^21^. At this junction, ICD and ICL compete for the common substrate, isocitrate. ICD and ICL are each present as two isoforms in most clinical *M. tuberculosis* strains^22^, but some clinical strains, including the commonly used H37Rv laboratory strain, carry a frameshift mutation in the *icl2* gene that generates two truncated gene products lacking detectable ICL activity^23^. The relative activities of *M. tuberculosis* ICD and ICL are controlled by multiple regulatory layers. At the transcriptional level, ICL1 and ICL2 have been reported to be regulated by different transcription factors under varying nutritional conditions^24–28^. Metabolite-mediated regulation has also been described^13,19,21^, including the activation of ICL2 by acetyl-CoA^29–30^, which accumulates during growth on fatty acids. Cross-activation of ICD2 from related species *M. smegmatis* by ICL-derived glyoxylate has also been reported^19^. Post-translational modifications, particularly lysine acylation, have further been proposed to regulate ICD and ICL^31–33^. For example, ICD1 was reported to be inhibited via lysine acetylation by the acetyltransferase Rv2170^33^, although this effect could not be reproduced in our laboratory (unpublished). However, how these regulatory inputs work together to dynamically coordinate carbon flux partitioning at the TCA cycle-glyoxylate shunt junction in response to carbon nutrient availability remains unclear.

Here, we combine gene expression and metabolomics in *M. tuberculosis* with biochemical approaches using recombinant enzymes to investigate how the bacterium responds to nutrient changes. We have developed a novel NMR-based in vitro assay that reconstitutes this metabolic junction at defined enzyme and metabolite concentrations to quantify carbon flux under defined conditions. This enables approximation of the intracellular environment under physiologically relevant conditions and allows us to define the molecular mechanisms governing carbon flux at this central metabolic junction. We show that only *icl1* expression responds significantly to changes in carbon substrate availability, whereas *icd1*, *icd2* and *icl2* do not. In addition, we uncover extensive regulation of ICDs and ICLs by metabolite signals from upstream, downstream and connected pathways, including the activation of ICD2 by glycolytic metabolites and TCA cycle intermediates associated with entry into gluconeogenesis. Importantly, integration of these regulatory layers reveals a net flux behaviour that cannot be inferred when individual layers are examined in isolation. Together, these analyses reveal a ‘mixer-tap’ mechanism that dynamically partitions carbon flux between the TCA cycle and the glyoxylate shunt to balance energy production with carbon conservation.

## Results

### *M. tuberculosis icl1* gene expression responds selectively to changes in carbon substrate

To determine how *M. tuberculosis* ICD and ICL are differentially regulated by carbon source availability, we first measured the expression of the ICD and ICL isoforms in *M. tuberculosis* grown in glucose- or acetate-containing media. In contrast to previous studies that examined expression under steady-state growth on a single carbon source^5,18^, our analyses focused on time-dependent responses following a switch from glucose to acetate (and vice versa) (**Fig. 1b**). This approach allowed us to capture how quickly *M. tuberculosis* shifts the expression of its ICD and ICL isoforms in response to carbon source changes over time.

To perform the carbon switch experiment, *M. tuberculosis* CDC1551 was first cultured in media containing one carbon substrate (substrate of origin) until an OD_600_ of ∼0.4 was reached. The bacteria were then pelleted and resuspended in fresh media containing the new carbon substrate (substrate of switch) (**Fig. 1b**). CDC1551 was used, as it is a common laboratory strain known to encode the full-length, catalytically competent ICL2^34^, unlike H37Rv^23^. Bacterial samples were collected at four timepoints (0, 3, 16, 72 hours) after the carbon switch, and the expression of *icd1*, *icd2*, *icl1* and *icl2* were then analysed.

Transcripts of all isoforms were detected under all culture conditions (**Extended Data Fig. 1**). Among all the changes observed, only *icl1* showed significant expression changes in response to carbon source switching at all timepoints after the media switch (**Fig. 1c**). *icl1* was significantly upregulated following a switch from glucose to acetate and significantly downregulated following a switch from acetate to glucose. These are consistent with prior reports that ICL1 protein abundance and activity are induced during growth on non-glycolytic carbon sources and reflect the requirement for glyoxylate shunt engagement under these conditions^19,21^. Most notably, this response was detected as early as the first timepoint measured, 3 hours after the media switch, indicating rapid transcriptional adaptation to the new carbon substrate.

### *M. tuberculosis* ICD and ICL activities are extensively modulated by central carbon metabolites

The limited transcriptional regulation at the TCA cycle-glyoxylate shunt junction prompted us to examine whether *M. tuberculosis* ICD and ICL are regulated by other mechanisms. Given that central carbon metabolite concentrations are shaped by carbon substrate availability and utilisation, and that prior studies have identified that the activities of some ICD and ICL isoforms are influenced by metabolites^19,21, 29^, we next focused on building a comprehensive picture of protein-metabolite interactions. We began by systematically screening metabolites from glycolysis, the TCA cycle, and the glyoxylate shunt for their effects on the activity of *M. tuberculosis* ICD1/2 and ICL1/2. Initial screens were conducted using individual enzyme isoforms with a single metabolite at a concentration of 5 mM. To minimise false positives, the concentration of isocitrate (substrate) was optimised such that a competitive inhibitor with a *K*_i_ of 1 mM would produce at least 50% inhibition at 5 mM concentration (**Supplementary Equations**; **Supplementary Table 1**).

Our screening shows that ICD and ICL are extensively regulated by metabolites and reveals a complex pattern of control (**Fig. 2a**). Both metabolite-mediated inhibition and activation of ICD and ICL were observed, with three metabolites (pyruvate, glyoxylate and oxaloacetate) appearing to exert opposing effects on the activities of the two enzymes, activating ICD2 and inhibiting ICL1/2 (**Fig. 2a**). Moreover, reaction products, including NADPH/2-oxoglutarate for ICD and glyoxylate/succinate for ICL, were found to differentially modulate enzyme activity by acting as product inhibitors, cross-activators or cross-inhibitors (**Fig. 2a**). These results indicate that ICD and ICL activity are likely regulated in an interdependent and coordinated manner.

**Figure 2.**
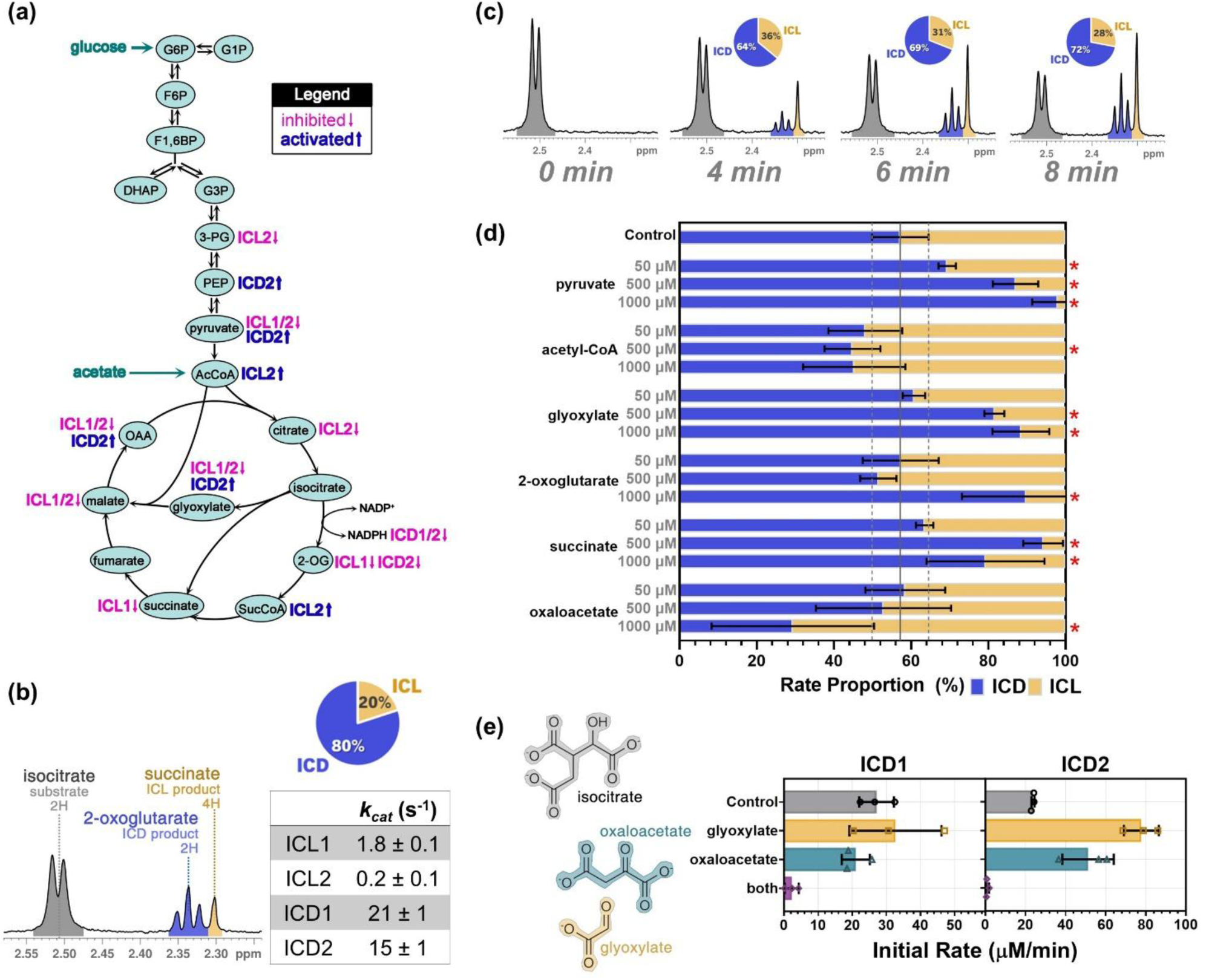
Metabolite effectors of *M. tuberculosis* ICD and ICL. **(a)** Effect of central carbon metabolites on *M. tuberculosis* ICL1/2 and ICD1/2 activity. Glycolytic and TCA cycle metabolites are mapped onto their respective pathways. Effects greater than 50% are indicated for the relevant isoform. Blue indicates activation, and pink indicates inhibition. Metabolite concentration was 5 mM. Reactions were performed at 25 °C and contained enzyme, _DL_-isocitrate, 5 mM metabolite, and 5 mM MgCl_2_. See **Supplementary Table 1** for substrate and enzyme concentrations. Refer to **Supplementary** Fig. 1 for the raw data on ICD and ICL reaction rates. For ^1^H NMR-based kinetics, 50 mM Tris-d_11_ in 90% H_2_O/10% D_2_O was used. For absorbance-based kinetics, 10 mM phenylhydrazine was added for ICL reactions, and 50 mM Tris (pH 7.5) was used as the buffer. **(b)** ^1^H NMR allows the simultaneous monitoring of ICD- and ICL-catalysed reactions. The isocitrate (2.51 ppm; 2H; common substrate), 2-oxoglutarate (2.34 ppm; 2H; ICD product), and succinate (2.30 ppm; 4H; ICL product) peaks were used to quantify the ICL and ICD reaction rates. The spectra shown was measured 3.5 min after enzyme addition. The relative proportion of the ICL (yellow) and ICD (blue) is represented as a pie chart. Reaction mixture contained 37.5 nM of each of the ICL and ICD isoforms, 1 mM _DL_-isocitrate, 1 mM NADP^+^, 5 mM MgCl_2_, 0.02% NaN_3_, 50 mM Tris-d_11_ in 90% H_2_O and 10% D_2_O. Catalytic efficiencies of the ICL1/2 and ICD1/2 are shown in a table format. See **Supplementary Table 2** for full kinetic parameters. **(c)** A time-dependent increase in the ICD reaction rate relative to the ICL reaction rate was observed. The reaction conditions are given in (b). **(d)** The relative proportions of ICD and ICL reaction rates at different metabolite concentrations are shown. Only metabolites that produced proportions significantly different from the control are included in this figure. Refer to **Extended Data Fig. 3** for the full data including all the metabolites tested. Asterisks (*) indicate statistically significant differences (p < 0.05) relative to the no metabolite control using the Student’s t-test corrected for multiple comparisons using the Holm-Sidak method. Error bars represent the s.d. from three replicates. Reaction mixture contained 12.75 nM of each ICL, 2.25 nM of each ICD, 200 mM _DL_-isocitrate, 1 mM NADP^+^, 5 mM MgCl_2_, 0.02% NaN_3_, 50 mM Tris-d_11_ in 90% H_2_O and 10% D_2_O. **(e)** The effect of glyoxylate-oxaloacetate mixture on the activity of *M. tuberculosis* ICD1 and ICD2. Error bars represent s.d. from three replicates. The reactions were measured at 25 °C using the absorbance-based assay. The mixtures contained 25 nM ICD1 / 50 nM ICD2, 250 µM _DL_-isocitrate, 100 µM (ICD1) / 1 mM (ICD2) NADP^+^, 5 mM MgCl_2_ in 50 mM Tris (pH 7.5). 5 mM of each metabolite or a 5 mM mixture (2.5 mM glyoxylate + 2.5 mM oxaloacetate) was added where applicable.

In the case of ICD, metabolite-mediated regulation was largely confined to ICD2. We found that the activity of *M. tuberculosis* ICD2 was activated by glyoxylate, in agreement with the previous study that showed glyoxylate-activation of *M. smegmatis* ICD2^19^. Our screening additionally uncovered three novel activators of ICD2: pyruvate, oxaloacetate, and phosphoenolpyruvate (**Fig. 2a**). Pyruvate and phosphoenolpyruvate are intermediates of glycolysis, while oxaloacetate is a gluconeogenic precursor. Dose response curves show that pyruvate, oxaloacetate and glyoxylate can increase the activity of ICD2 by up to 4 times (**Extended Data Fig. 2b**), and Michaelis-Menten analyses reveal allosteric activation (no change in *K*_M_, increase in *V*_max_) (**Supplementary Fig. 2**). Phosphoenolpyruvate, however, only showed up to a 2-fold increase in activity and only at concentrations above 3 mM (**Extended Data Fig. 2b**). Phosphoenolpyruvate can be hydrolysed to pyruvate^35^, and the ^1^H NMR spectrum of our phosphoenolpyruvate stock solution showed a peak corresponding to pyruvate (**Supplementary Fig. 3**). Thus, the modest activation observed with phosphoenolpyruvate may in part arise from break down to pyruvate, though we cannot exclude a direct activating effect of phosphoenolpyruvate itself.

In contrast, both ICL isoforms were heavily influenced by metabolites. We found that glyoxylate, pyruvate and oxaloacetate, in addition to their roles as activators of ICD2, were also inhibitors of ICL1 and ICL2 (**Fig. 2a**). ICL1 and/or ICL2 are also inhibited by 3-phosphoglycerate (glycolysis intermediate) and malate (gluconeogenic precursor). Binding studies showed that these metabolite inhibitors are competitive against substrate binding (**Supplementary Fig. 2a**). Together with our prior findings that ICL2 is activated by the fatty acid metabolite acetyl-CoA^29,30^, these results indicate that carbon flux through the glyoxylate shunt can be either enhanced or suppressed depending on the presence of fatty acid- or glycolysis-derived metabolites. This is consistent with the canonical substrate-dependent role of the glyoxylate shunt.

### Competitive NMR analysis reveals complexity in metabolite-mediated regulation of *M. tuberculosis* ICD and ICL

Our screening experiments were conducted at a single high metabolite concentration (5 mM), which may not reflect physiological levels, so we performed metabolomic experiments to estimate the intracellular concentration range of the effectors. Using *M. tuberculosis* CDC1551 and following the same substrate switch experimental design as the transcriptional analysis (**Fig. 1b**), the concentration of 12 metabolites (2-oxoglutarate, citrate, isocitrate, aspartate, malate, fumarate, CoA, succinate, phosphoenolpyruvate, acetyl-CoA, 3-phosphoglycerate and pyruvate) were measured and quantified this time at 0, 2, 16 and 72 hours. The second timepoint was measured at 2 hrs instead of 3 hrs (used for the gene expression experiments) to see whether metabolite changes occurred before gene expression changes. Data indicate that the cellular concentrations of metabolite effectors typically fell within the 0–1 mM range, while intracellular isocitrate levels were around 25–55 μM (**Supplementary Fig. 4a**). We were unable to accurately quantify the intracellular concentration of glyoxylate due to its low molecular weight, volatility and high reactivity^36^. Unsupervised hierarchical clustering and correlation analysis of our metabolomics dataset revealed four distinct clusters based on the metabolites’ patterns of change (**Supplementary Fig. 4b**). This suggests that the different groups of metabolites fluctuate differently according to nutrient availability.

We then investigated whether metabolic effectors influence ICD and ICL activities at physiologically relevant concentrations. Rather than screening individual isoforms in isolation, we studied all ICD and ICL isoforms together to understand how their activities are partitioned in a coordinated manner by metabolite effectors. We applied ^1^H NMR to simultaneously monitor both reactions, as this method allows separate peaks for isocitrate (common substrate), 2-oxoglutarate (ICD product) and succinate (ICL product) to be resolved and quantified (**Fig. 2b**; Refer to **Supplementary Fig. 5** for the full spectrum) ^37,38^.

We first performed an NMR time course experiment using equal amounts of all four enzymes (ICD1, ICD2, ICL1 and ICL2) to validate the methodology. Michaelis-Menten kinetics indicate that ICD1 and ICD2 have a higher catalytic efficiency than ICL1 and ICL2 (**Fig. 2b**). Consistent with this, in our NMR time course, the intensity of the 2-oxoglutarate resonance (ICD product) increased more rapidly than that of succinate (ICL product) (**Fig. 2b**). Notably, a time-dependent increase in the ICD-to-ICL reaction rate ratio was observed (**Fig. 2c**). This could arise from cross-activation of ICD2 by ICL-generated glyoxylate and/or cross-inhibition of ICL1 by ICD-generated 2-oxoglutarate. To distinguish between these possibilities, we repeated the time course experiments using both ICL1 and ICL2 but only a single ICD isoform, as glyoxylate activates ICD2 but not ICD1. Under these conditions, the time-dependent increase in the ICD-to-ICL reaction rate ratio was observed with ICD2 but not with ICD1 (**Extended Data Fig. 2c**), showing that the activation of ICD2 by glyoxylate dominates over inhibition of ICL by 2-oxoglutarate. This also highlights the highly dynamic nature of the relative activities between ICD and ICL.

We next applied the NMR assay to examine how the activities of ICD and ICL are jointly modulated by metabolites. Individual metabolites were screened at three physiologically relevant concentrations (50, 500 and 1000 µM) were tested. For the substrate, 200 mM DL-isocitrate (corresponding to 100 μM of the physiologically relevant D-isomer) was used. The amounts of ICD and ICL (12.75 nM each of ICL1 and ICL2, and 2.25 nM each of ICD1 and ICD2) was also optimised so that the proportion of ICD and ICL initial reaction rates were approximately 50%-50% in the absence of any metabolite effectors to aid the visualisation of metabolite-mediated changes.

Among the metabolite effectors identified from our initial screen, six metabolites (pyruvate, glyoxylate, 2-oxoglutarate, succinate, oxaloacetate and acetyl-CoA) significantly altered the ICD-to-ICL reaction rate ratio compared with the control (**Fig. 2d**). In particular, pyruvate, oxaloacetate, and succinate altered the ICD-ICL reaction proportion by more than 15% at their respective physiological concentrations (**Fig. 2d**; **Supplementary Fig. 4a**). Consistent with their individual regulatory effects, pyruvate and glyoxylate (ICD2 activators and ICL1/2 inhibitors), as well as 2-oxoglutarate (ICL1/2 inhibitor) and succinate (ICL1 inhibitor), increased the proportion of ICD reaction relative to ICL in the NMR assay (**Fig. 2d**). Similarly, acetyl-CoA, an activator of ICL2, increased the proportion of ICL reaction relative to ICD (**Fig. 2d**). In contrast, oxaloacetate, which is an activator of ICD2 and an inhibitor of ICL1 and ICL2 (**Fig. 2a**), was found to promote the relative ICL activity over ICD in the NMR assay (**Fig. 2d**). Examination of the absolute reaction rates revealed that oxaloacetate inhibited both ICL and ICD in the mixed reaction, with the inhibitory effect being stronger for ICD (**Extended** Data Fig. 3).

As our assay contained both ICD and ICL, products of both ICD- and ICL-catalysed reactions were formed during the NMR assay. It is therefore possible that these reaction products may alter the effect of oxaloacetate on ICD2. The inhibition of ICD by a glyoxylate-oxaloacetate mixture has been reported from different organisms^39–43^. We therefore tested the effects of oxaloacetate in the presence of glyoxylate on ICD. Consistent with this hypothesis, we found that glyoxylate and oxaloacetate (which are both activators of ICD2 on their own) as a mixture inhibited both ICD1 and ICD2, and that this inhibition was competitive with respect to isocitrate (**Fig. 2e**; **Supplementary Fig. 2**). This effect may arise from the structural similarity of glyoxylate and oxaloacetate to isocitrate (**Fig. 2e**). These observations indicate that metabolite regulation is unlikely to be driven by single effectors in isolation, but instead from the combined actions of multiple metabolites.

### Integrated gene expression and metabolite-driven activity modulation reveal dynamic regulation at the TCA cycle-glyoxylate shunt junction

Our results suggest that the activities of ICD and ICL are shaped by both gene expression (and thus enzyme abundance) and metabolite regulation. However, how these layers of control interact to modulate flux at the ICD-ICL junction remains unclear, as we show that metabolites exert opposing and synergistic effects that cannot be predicted from individual measurements. We therefore applied the ¹H NMR assay to examine how carbon flux is partitioned at this metabolic branchpoint using the gene expression and metabolomics data derived from our carbon switch experiments.

We first estimated the relative abundance of the ICD and ICL isoforms using our time-resolved gene expression data (**Extended Data Fig. 1**; **Supplementary Table 3**). Although gene expression does not directly reflect protein abundance, our data showed that only *icl1* expression changed significantly in response to carbon substrate switching (**Fig. 1c**). This is consistent with previous observations that ICL1 protein abundance increases during growth on acetate, while ICL2, ICD1 and ICD2 remain largely unchanged^19^, supporting the use of expression data as a reasonable proxy for relative isoform abundance under these conditions.

For metabolite perturbation, we selected a mixture of 11 effectors based on our screening data and quantitative targeted metabolomics measurements in *M. tuberculosis* under carbon switch and control conditions (**Supplementary Table 4**). In the absence of any metabolite mixtures, the reaction rates correlated with enzyme concentration and their intrinsic kinetics (**Fig. 3a**). When the metabolite mixtures (**Supplementary Table 5**) were added, the absolute ICD reaction rates generally increased, while the absolute ICL reaction rates decreased (**Fig. 3b**).

**Figure 3.**
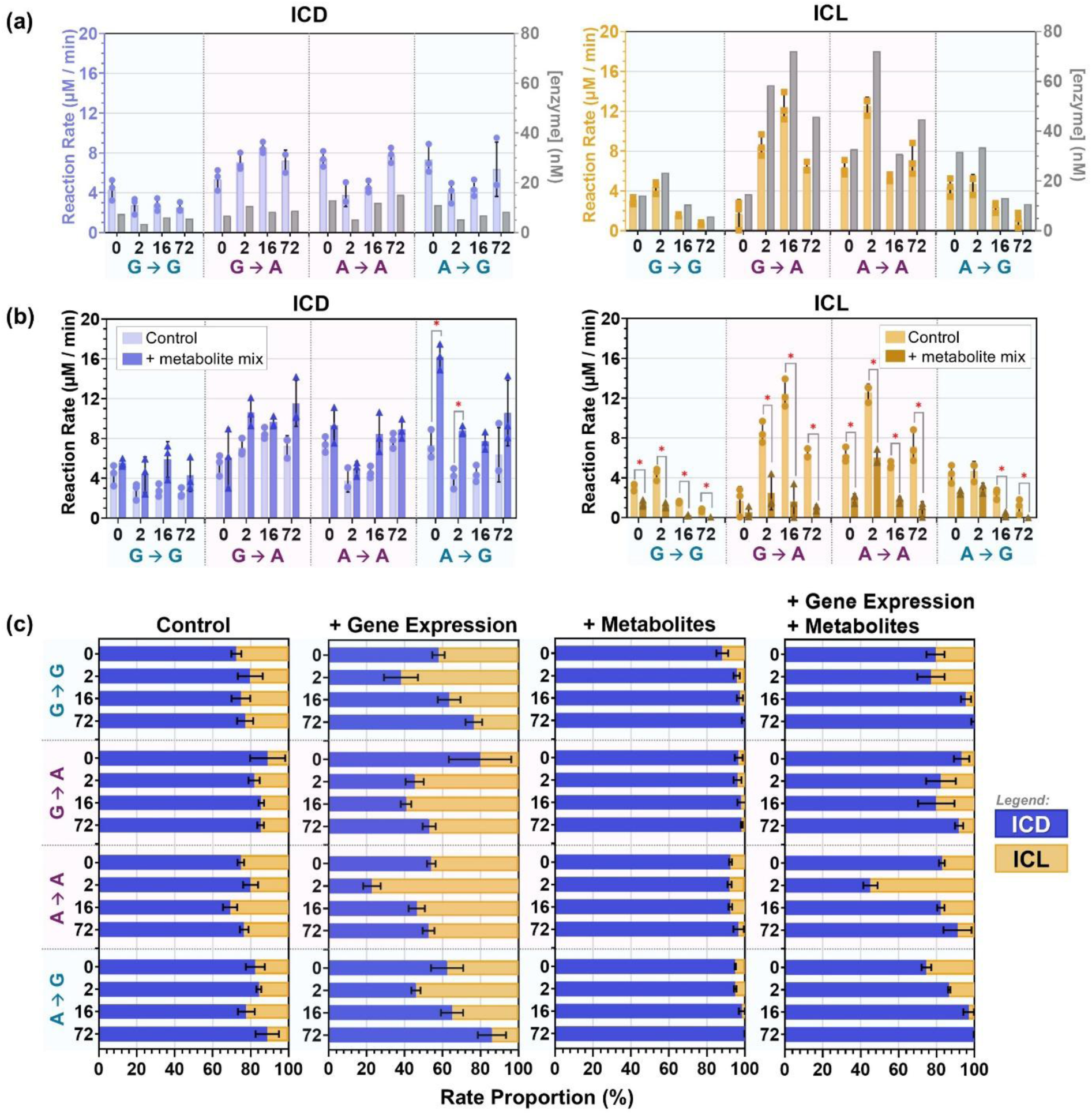
Incorporating gene expression and metabolomics data into the ^1^H NMR model of the ICL-ICD junction. Error bars represent s.d. from three replicates. G represents the condition where *M. tuberculosis* was grown in media containing glucose, and A represents the condition where *M. tuberculosis* was grown in acetate. The arrow indicates the switch in substrate (i.e. G ➔ A means *M. tuberculosis* grown in glucose then switched to fresh media containing acetate). The enzyme concentration ratios were determined from absolute gene expression data (**Supplementary Table 3**), while the metabolite effector concentrations were determined from the approximate intracellular concentrations obtained from our metabolomics data (**Supplementary Table 4**). **(a)** Effect of enzyme concentration on reaction rates within the ^1^H NMR model of the ICL-ICD junction. The reaction rates for ICD (left) and ICL (right) are shown in blue (ICD) or yellow (ICL) bars and plotted on the left y-axis. The enzyme concentrations used were plotted in grey bars on the right y-axis. **(b)** The effect of metabolite mixtures on the ICD (left) and ICL (right) rate within the ^1^H NMR model of the ICL-ICD junction. The reaction rates of ICD and ICL were measured simultaneously in the presence or absence of the metabolite mixtures. **(c)** The effect of enzyme abundance and metabolite mixtures on the proportion of ICD and ICL rates. The proportions of ICD (blue) and ICL (yellow) were calculated by expressing their respective reaction rates as a percentage of the total ICD and ICL reaction rates. For the ‘Control’ and ‘+ Metabolites’, the effect of enzyme abundance was eliminated by normalising the ICD and ICL rates using the ICD and ICL enzyme concentrations, respectively. The rates were then expressed as a percentage of the total ICD and ICL normalised reaction rates.

In contrast, the relative reaction rates of ICD and ICL, and therefore flux partitioning between the TCA cycle and the glyoxylate shunt, were more complex (**Fig. 3c**). A dissection of the factors revealed that gene expression favoured an increase in the ICL reaction rate. This is in accordance with our gene expression studies where only *icl1* gene expression showed significant responses to a change in carbon substrate (**Fig. 1c**). On the other hand, the metabolite mixture showed almost a complete diversion towards the ICD reaction, consistent with our metabolite screening results where most of the effectors were either ICD2 activators or ICL1/2 inhibitors (**Fig. 2a**; **Fig. 3b**). These two regulatory mechanisms balanced each other out to generate the final flux that varied according to time.

This dynamic redistribution differs from the binary on-off regulation of the glyoxylate shunt described in several other bacterial systems^16^, or even a single rheostat as previously proposed for mycobacteria^19^. Instead, our results suggest a more adaptive mode of flux control at this metabolic junction in *M. tuberculosis*, more analogous to a mixer-tap, where water flow rate and temperature mixing are carefully balanced, as are enzyme abundances and metabolite combinations (**Fig. 4a**). A close investigation into these regulatory factors at different timepoints explains the ICD-ICL rate partitioning under the different culture conditions (**Fig. 4b**).

**Figure 4.**
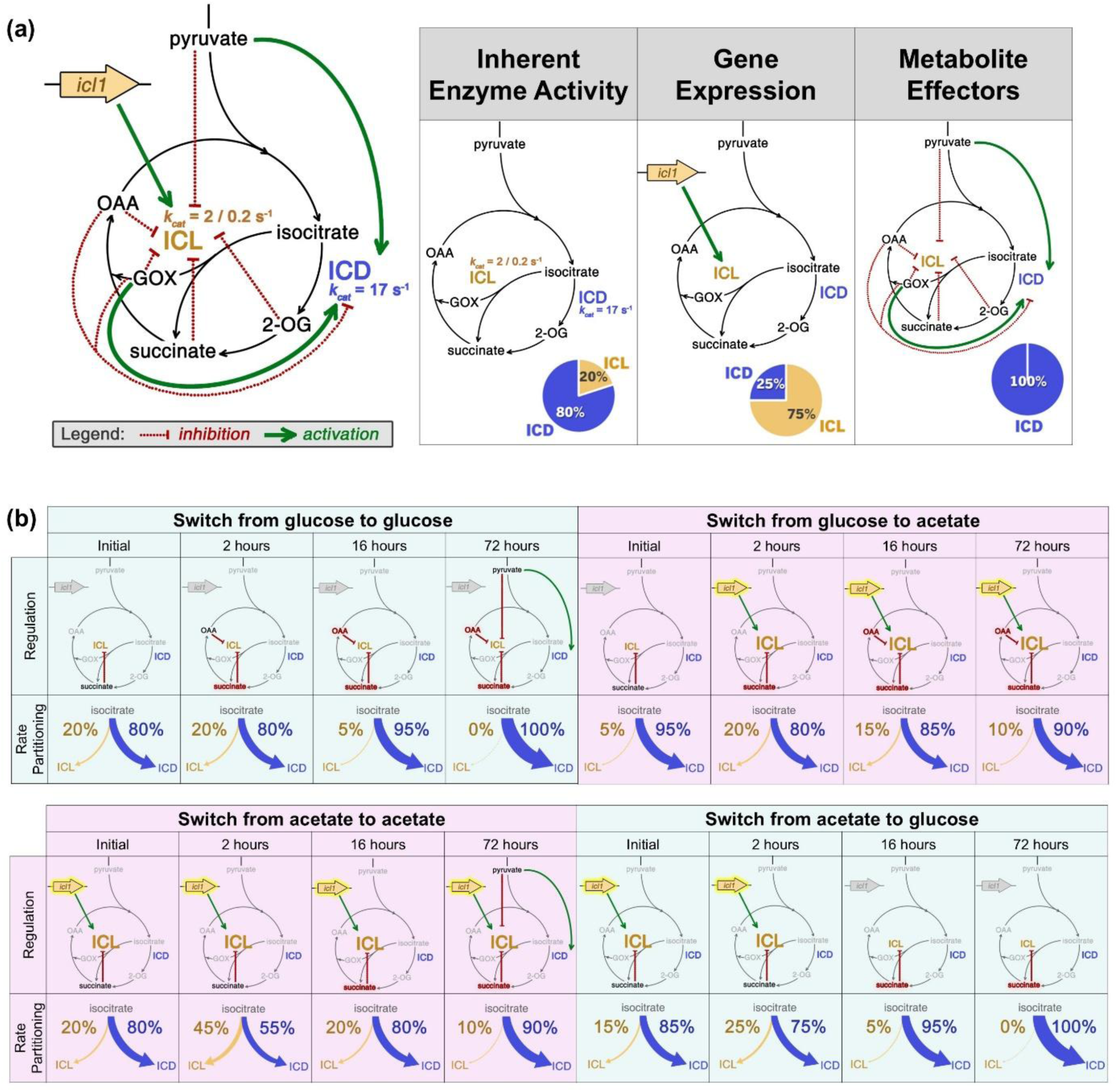
Model of the regulatory mechanisms governing the TCA cycle-glyoxylate shunt junction in *M. tuberculosis*. **(a)** The junction is regulated by a combination of inherent enzyme activity, gene expression, and metabolite inhibition/activation. Red dotted lines represent inhibition, and green solid lines represent activation. Abbreviations: GOX = glyoxylate; 2-OG = 2-oxoglutarate; OAA = oxaloacetate. The most extreme rate proportion between ICL (blue) and ICD (yellow) measured in the *in vitro* model is illustrated in pie charts. **(b)** The different regulatory mechanisms involved in *M. tuberculosis* during the carbon switch experiment. The *icl1* gene expression is upregulated when acetate is introduced, as indicated by the font size of ‘ICL’. If the intracellular concentration can affect the rate proportion of ICL and ICD by 15% (based on the approximate EC_50_ curves in **Extended Data Fig. 3b**), the metabolite font is coloured in black. If the proportion can be changed by more than 25%, the metabolite font is coloured in red. The corresponding rate proportions from the different culture conditions are taken from Fig. 3c.

When *M. tuberculosis* was grown on glucose (glucose-to-glucose control), we observed a progressive increase in ICD activity over time, reaching ∼100% by 72 hours (**Fig. 4c**). At early time points, most metabolite effectors were not present at physiological concentrations sufficient to activate or inhibit ICD or ICL, and the relative flux at the junction therefore reflected the inherent catalytic activities of the enzymes. However, as the bacteria continued to grow on glucose, the concentrations of key metabolites increased, including pyruvate (an ICD2 activator and ICL1/2 inhibitor), succinate (an ICL1 inhibitor) and oxaloacetate (an ICD2 activator and ICL1/2 inhibitor) (**Fig. 4c**; **Supplementary Fig. 4a**). The progressive redirection of flux towards the TCA cycle is therefore likely driven by the combined activation of ICD and inhibition of ICL by these accumulating metabolites.

In contrast, although the acetate-to-glucose transition displayed a similar overall trajectory (converging on ∼100% ICD activity over 72 hours) (**Fig. 4c**), it was regulated by a different mechanism. At early time points following the carbon switch, ICL1 remained highly expressed as the bacterium adapted from acetate metabolism (**Fig. 1c**; **Extended Data Fig. 1**). Under these conditions, ICL activity was partially offset by elevated succinate levels. As growth on glucose continued, ICL1 expression declined, coinciding with a further increase in succinate concentration (**Fig. 1c**; **Extended Data Fig. 1**; **Supplementary Fig. 4a**). Together, these effects resulted in progressive suppression of ICL activity, ultimately directing flux almost entirely through the TCA cycle.

*M. tuberculosis* maintained on acetate (acetate-to-acetate control) requires glucose synthesis to support biomass production. As a result, we observed upregulation of *icl1* over time, which diverted isocitrate turnover towards the glyoxylate shunt to preserve carbon for gluconeogenesis, resulting in an approximately balanced flux distribution between the glyoxylate shunt and the TCA cycle (**Fig. 4c**). We also observed a decrease in glycolytic intermediates, including pyruvate (**Supplementary Fig. 4a**). This is likely due to the cells consuming its glucose store. As growth on acetate continued, however, the concentrations of pyruvate (an ICD2 activator and ICL1/2 inhibitor) and succinate (an ICL1 inhibitor) increased (**Supplementary Fig. 4a**), counteracting the effect of *icl1* upregulation and progressively redirecting flux towards the TCA cycle. These results imply that transcriptional responses respond immediately to the presence of non-glycolytic substrates through *icl1* upregulation, while metabolite effectors act gradually on the ICD-ICL junction as the cell balances its requirements.

The rebound in concentration of glycolytic metabolites is consistent with glucose regeneration via gluconeogenesis, and the shift away from the glyoxylate shunt may reflect the reduced reliance on carbon conservation once gluconeogenic demands are met. A similar fluctuation, and tight balance between elevated ICL activity and metabolite effectors that oppose it, was observed in the glucose-to-acetate transition samples. Overall, the behaviour of the ICD-ICL junction under the different culture conditions is consistent with the dual metabolic demands imposed by acetate or glucose, where carbon must be partitioned between energy generation via the TCA cycle and conservation for biosynthesis through gluconeogenesis.

## Discussion

The TCA cycle-glyoxylate shunt junction is a tightly regulated and essential metabolic node in *M. tuberculosis* that adjusts flux partitioning under different substrate conditions to balance energy generation with carbon conservation for biosynthesis^4,6,19^. It is now established that regulatory mechanisms described in other bacterial systems are not directly applicable to *M. tuberculosis*^16^. In this study, by evaluating regulation at this node across multiple layers, we show that it is far more intricate than previously described. Not only have we identified novel metabolite effectors, but we also demonstrate that ICD and ICL activities are shaped by the combined actions of multiple previously unlinked metabolites, alongside differences in gene expression and enzyme abundance. As a result, the net effect on carbon flux distribution cannot be predicted *a priori* from any single experiment. To address this, we applied an information-rich NMR-based enzyme assay to examine how these factors interact to fine-tune regulation at the ICL-ICD junction. This work establishes a quantitative, systems-level framework linking gene expression, metabolite dynamics and enzyme activity to carbon flux partitioning at this central metabolic node. The experimental framework developed here captures the complexity of *in vivo* protein and metabolite mixtures while retaining the clarity of an *in vitro* system, providing a more integrated and quantitative view of regulation at this metabolic node.

Previous studies reported increased ICL activity during growth on acetate^19,21^, although the temporal dynamics were not resolved. In our experimental system, this increase in ICL activity was most evident under conditions where metabolite effector concentrations remained low. As these metabolites accumulated, their opposing effects on ICL activity became more pronounced, suggesting that they may act as indicators of sufficient anaplerotic replenishment, at which point reliance on the glyoxylate shunt diminishes. Together, these observations highlight the dynamic regulation of flux partitioning at this metabolic junction.

Our data show that changes in metabolite levels are critical in shaping flux at the ICL-ICD junction. Analysis of our metabolomics dataset allowed us to group these effectors into four distinct clusters based on their patterns of change. Notably, isocitrate, the shared substrate of ICD and ICL, together with three prominent modulators identified above (pyruvate, succinate and oxaloacetate), all fell into separate clusters. This highlights the difficulty of inferring net regulatory outcomes from any single metabolite alone, and supports the view that carbon substrate availability drives distinct metabolic states in *M. tuberculosis*, with the resulting metabolite landscape collectively regulating flux partitioning across different stages of growth and adaptation. Further studies are needed to understand how *M. tuberculosis* remodels its metabolic landscape in response to nutrient availability.

Collectively, these findings indicate that flux partitioning at the TCA cycle-glyoxylate shunt junction is governed by an integrated and highly dynamic regulatory framework at both the gene- and protein-level. Rather than being controlled by a single dominant factor, the relative activities of ICD and ICL emerge from the combined influences of metabolite effectors, enzyme abundance and broader metabolic state. These regulatory inputs act in a time-dependent manner, with shifting metabolite landscapes progressively reshaping enzyme activity and redirecting carbon flux as cells adapt to changing nutrient environments. This capacity for fine-tuned flux control may contribute to the ability of *M. tuberculosis* to persist under nutrient-limited and host-associated conditions.

In this context, the junction does not operate as a simple binary switch between the TCA cycle and the glyoxylate shunt. Instead, our data support a model in which flux is continuously redistributed in a graded manner, rather than governed by discrete regulatory switches, analogous to a metabolic ‘mixer tap’ where multiple regulatory inputs are integrated to fine-tune the balance between energy generation and carbon conservation. Such a mechanism allows *M. tuberculosis* to respond flexibly to fluctuating substrate availability and biosynthetic demands, enabling coordinated metabolic adaptation across different stages of growth and persistence.

To fully understand how carbon flux regulation influences *M. tuberculosis* survival, this approach can be extended to a broader range of conditions, including host-relevant models such as macrophage infection. Importantly, this integrative NMR-based framework provides a robust and extensible platform to interrogate this metabolic junction and its role in adaptation and survival, and can be readily expanded to incorporate additional pathways and regulatory inputs as they are identified. More broadly, this approach establishes a general framework for dissecting how multiple regulatory layers interact to control metabolic flux, with potential applicability to other metabolic systems beyond *M. tuberculosis*.

## Methods

### Materials

Unless otherwise stated, all chemicals were purchased from Sigma-Aldrich/Merck, ThermoFisher Scientific, AK Scientific, or Bio-Rad. Tris-d_11_ and D_2_O were from Cambridge Isotope Laboratories or Cortecnet. Competent cells were obtained from Agilent.

### Bacterial strains and culture conditions

All *M. tuberculosis* CDC1551 cultures were conducted as standing cultures in a 37 °C incubator with 5% CO_2_ inside a BSL3 facility. For growth on different carbon sources, a starter culture of CDC1551 was grown in 30 mL of 7H9 supplemented with OADC, 0.5% glycerol, and 0.05% Tween 80. 2.5 mL of the starter culture was inoculated into 125 mL 7H9 media supplemented with 0.5% BSA, 0.085% NaCl, 0.05% tyloxapol, and 0.2% glucose or acetate. This 7H9 minimal media (supplemented with BSA, NaCl, and tyloxapol) has been used as a base medium in previous metabolomics studies to assess the response of *M tuberculosis* to different carbon substrates^44–46^. After 12 days, the bacteria were pelleted at 4000 *xg* for 5 min at 4 °C then resuspended into the 125 mL of 7H9 minimal media containing either glucose or acetate. For the gene expression study, 25 mL samples were taken at 0, 3, 16, and 72 hours. For the metabolomics study, 25 mL samples were taken at 0, 2, 16, and 72 hours, and these samples were normalised to the same OD_600_ by diluting with 7H9 media with 0.05% tyloxapol. 25 mL of the diluted samples was then pelleted for metabolomics analysis.

### RNA extraction and cDNA generation

The cell pellet was washed with ice-cold PBS then resuspended in 500 µL TRIzol and lysed via bead beating at 3 cycles of 30 sec at 9000 rpm and 2 °C with 1 min rest in between cycles (Bertin Technologies Precellys Evolution Touch Homogeniser fitted with Cryolys Evolution). RNA was separated from TRIzol samples using Phasemaker tubes (Life Technologies) with addition of chloroform:isoamyl alcohol (49:1 v/v) and precipitated with isopropanol, sodium acetate and linear polyacrylamide (Life Technologies) at -20 °C overnight. Precipitated RNA was washed twice with 80% (v/v) ethanol before final resuspension in distilled H_2_O. 1 µg of RNA extracted from *M. tuberculosis* cells was reverse transcribed using the High Capacity cDNA synthesis kit (Applied Biosystems) and diluted 1:2 with TE buffer (Invitrogen Life Technologies) containing 1 ng/mL Herring Sperm DNA (Promega).

### Gene expression by absolute quantitative Real-Time Polymerase Chain Reaction (qRT-PCR)

Absolute qRT-PCR was performed with Fast SYBR Green (Applied Biosystems), using 10% cDNA in duplicate on QuantStudio 12K system (ThermoFisher), with 60 °C annealing temperature and 40 cycles. Melt curve analysis was performed to confirm amplicon specificity. Absolute quantification was carried out for all genes using synthesised primers (IDT Technologies) and standard curves generated by serial dilution of target amplicon-containing plasmids (pGEM-T easy, Promega), to cover up to 6 logs of amplicon copy number per microliter with absolute copy number normalized to housekeeping gene sigma factor A (SigA) copy number. Stable expression of SigA across all samples was confirmed by Cycle Threshold (C_T_) comparison. An increase/decrease in gene expression levels was calculated using the mean quantity, comparing all samples to the control and plotted as absolute copy number per nanogram (ng) of cDNA.

Primers used for this study are listed below.

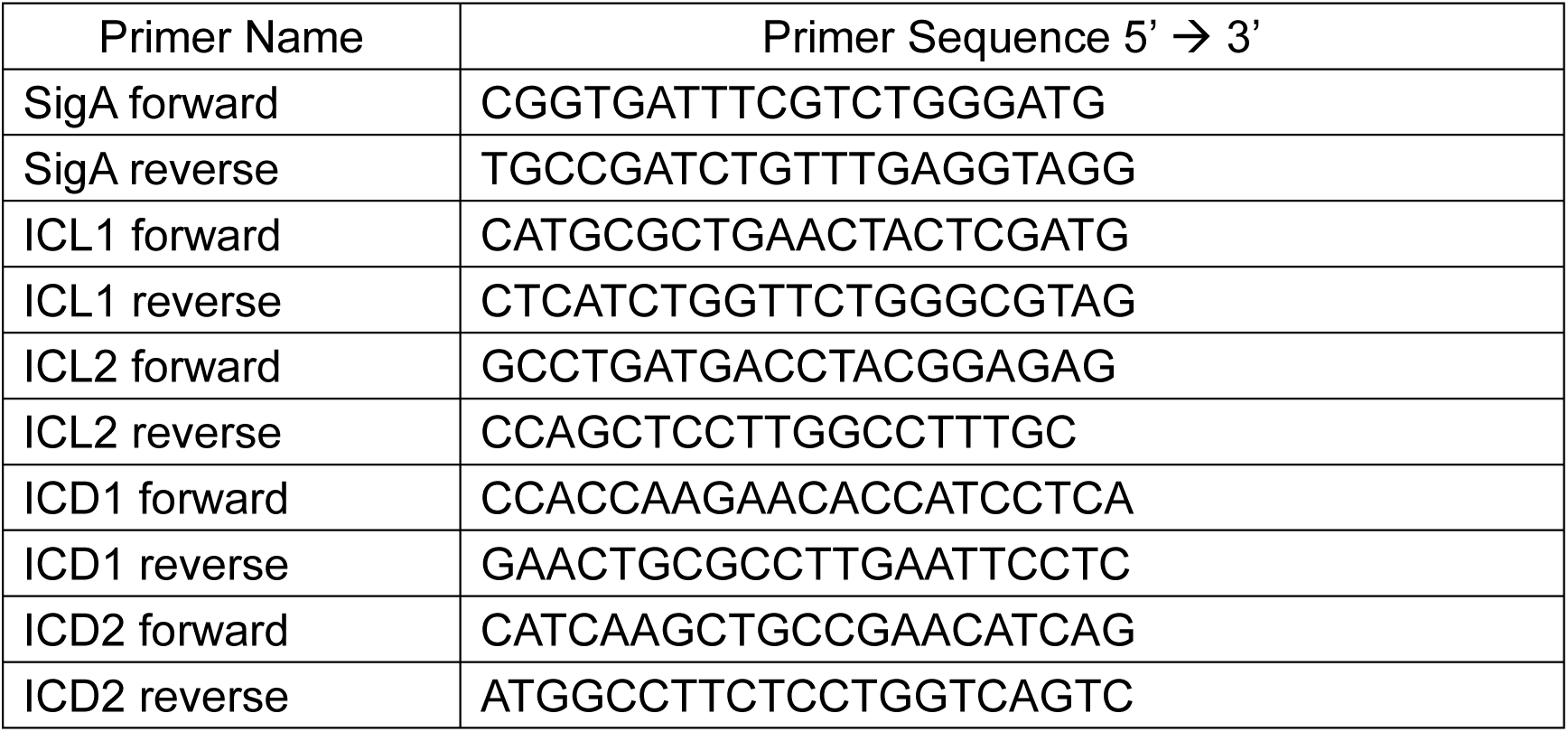

### Metabolite extraction

Endogenous concentrations of glycolysis and TCA intermediates (2-oxoglutarate, citrate, isocitrate, malate, fumarate, CoA, succinate, phosphoenolpyruvate, acetyl-CoA, 3-phosphoglycerate, and pyruvate) were measured by quantitative liquid chromatography-mass spectrometry (LC-MS). The cells were washed with ice-cold PBS then resuspended in 225 μL chloroform:methanol:water (2:6:1) and lysed by 6 cycles of flash freezing in liquid nitrogen, thawing, and vortexing. The tubes were centrifuged at maximum speed for 10 min at 4 °C, then the supernatants were transferred to a fresh tube for LC-MS analysis. 40 µL from each sample were pooled together to create pooled samples for the standard curves and for pooled quality control (PQC). Standard curves were generated by spiking the pooled samples with varying concentrations (1 nM to 100 μM) of chemical standards.

### Liquid chromatography-mass spectrometry (LC-MS)

Sample extracts were analysed by hydrophilic interaction liquid chromatography (HILIC) coupled to high-resolution mass spectrometry. Briefly, a polymeric iHILIC®-(P) Classic (HILIC Column, PEEK,150x4.6mm, 5µm, 200Å) at 25 °C with a gradient elution of 20 mM ammonium carbonate (Solvent A) and acetonitrile (Solvent B) (linear gradient time-%B as follows: 0 min-80%, 15 min-50%, 18 min-5%, 21 min-5%, 24 min-80%, 32 min-80%) on a Vanquish Horizon LC (Thermo Scientific). The flow rate was maintained at 500 μL/min. Samples were kept at 6 °C in the autosampler and 10 μL injected for analysis. Mass spectrometry was performed on a Q-Exactive Plus Orbitrap (Thermo Fisher Scientific, Australia) in MS1 polarity switching mode. The instrument was operated at 70,000 resolution, with the following conditions: Electrospray ionization voltage was 4 kV in positive and -3.5 kV in negative mode, capillary temperature = 300 °C; sheath gas = 50; Aux gas = 20; sweep gas = 2; probe temp = 120 °C). All samples were randomized for analysis and with PQC samples analysed every 10^th^ sample and blank injections before and after the run to assess instrument performance.

### Metabolomics data analysis

Peaks for targeted metabolites and associated standards were integrated in TraceFinder 4.1 (Thermo Fisher Scientific). For quantification, glycolysis and TCA intermediate concentrations were inferred from standard curves, using a linear equation with intercept set to zero (to remove endogenous concentration in the standard matrix) (i.e., area/linear coefficient = conc (µM)). The standard curves were made from six points (0.001, 0.1, 1, 10, 50, 100 µM) and generally showed good linearity with an R^2^ of 0.98 or above, with the exception of isocitric acid where we observe a slight deviation from linearity (R^2^ = 0.96). The measured metabolite concentrations in the extracts fell within the concentration range of the standard curves.

### Estimation of intracellular metabolite concentration

The volume of a tubercle bacillus was estimated according to a previous study^47^. It was assumed that one cell represents a cylinder with hemispherical ends with a width of 0.6 µm and length of 4 µm, giving a volume of 1.24 x 10^-15^ L per cell. Total cell numbers were calculated by using 25-mL cultures at 0.4 OD with the relationship that 1 OD unit corresponds to 5 x 10^8^ bacilli. This gives 6.2 µL cells per extract, and since this was diluted with 225 µL extraction solvent, this gives a dilution factor of 38.5. All the calculated metabolite concentrations from the metabolomics study were multiplied by this dilution factor to reflect the approximate intracellular concentration. The correlation between the metabolite concentrations over time under different culture conditions were analysed using MetaboAnalyst 6.0^48^.

### Recombinant protein production

Synthetic DNA fragments encoding ICL1 (*M. tuberculosis* H37Rv) and ICL2 (*M. tuberculosis* CDC1551) were obtained from Integrated DNA Technologies. The ICL1 gene was subcloned into pNIC28-Bsa4^49^, while the ICL2 gene was subcloned into pYUB28b^50^. The pNIC28-Bsa4 vector was a gift from Opher Gileadi (Addgene plasmid #26103). pET28a(+) plasmids encoding genes for ICD1 and ICD2 (*M. tuberculosis* H37Rv) were synthesised by GenScript. All proteins were synthesised with an N-terminal 6xHis tag.

Recombinant proteins were expressed in either *Escherichia coli* BL21 (DE3) cells or *E. coli* BL21 LOBSTR cells transformed with the pGro7 plasmid (Takara Bio Inc.) expressing the GroEL/GroES chaperones under the araB promoter. IPTG was added when the cell density reached an OD_600_ of 0.6-0.8. For proteins expressed with the GroEL/GroES chaperones, L-arabinose (final concentration of 0.1% w/v) was added when the cell density reached an OD_600_ of 0.3-0.4. The cells were harvested via centrifugation after overnight incubation at the target induction temperature. The proteins were purified by immobilised metal affinity chromatography (IMAC) and size exclusion chromatography. Purified aliquots were flash-frozen and stored at -80 °C until use. The expression condition and buffer systems used for each protein are given in **Supplementary Table 5** and **Supplementary Table 6**.

### Enzyme activity

The enzyme activities were measured either using the UV-visible absorbance-based assay or via NMR. Unless otherwise stated, the UV-visible absorbance-based assay was used. For the metabolite screening, NMR was used when the metabolite absorbance interfered with the product absorbance. Initial rates were calculated using data points up to 20% substrate turnover. To obtain the Michaelis-Menten or EC_50_ parameters, the initial rates were fitted to the Michaelis-Menten equation or the dose response curve using GraphPad Prism 8.

The UV-visible absorbance-based assays were conducted in 96-well plates with a final volume of 100 μL. For ICL, 10 mM phenylhydrazine was added to convert the glyoxylate product into a phenylhydrazone that absorbs light at 324 nm^51^. For ICD, the NADPH product absorbance was measured at 340 nm^52^.

NMR data was collected at 298 K using a Bruker 700 MHz AVANCE III HD equipped with a cryoprobe a Bruker 500 MHz AVANCE NEO. 5 mm diameter NMR tubes with a sample volume of 550 µL were used in all experiments. Solutions were buffered using 50 mM Tris-d_11_ (pH 7.5) dissolved in 90% H_2_O and 10% D_2_O. The pulse tip-angle calibration using the single-pulse nutation method (Bruker ‘pulsecal’ routine) was undertaken for each sample^53^. Time course experiments were performed based on previous studies^38^. The reactions were monitored by standard Bruker ^1^H experiments with water suppression by excitation sculpting (pulse program ‘zgesgp’)^54^. The number of transients was 16, and the relaxation delay was 2 sec. The lag time between the addition of the enzyme and the end of the first experiment was around 4 min. For experiments with a mixture of metabolite effectors, all spectra were subtracted by a reference spectrum of the metabolite mixtures alone in the absence of isocitrate and NADP^+^ before analysis.

## Acknowledgements

We would like to acknowledge the use of the Melbourne Magnetic Resonance, Melbourne Protein Characterisation, and Mass Spectrometry and Proteomics platforms at the Bio21 Molecular Biology and Biotechnology Institute, as well as the PC3 facility at the Walter and Eliza Hall Institute for this work. We would also like to thank Dr Rhys Grinter for his help in initiating the collaboration between the University of Melbourne and the Monash Proteomics & Metabolomics Platform. We also thank Dr Tianyang Liu for her help with sample preparation and useful discussions.

E.Y.W.H. is supported by the Melbourne Research Scholarship, Rowden White Scholarship, Norma Hilda Scholarship, Dame Margaret Blackwood Soroptimist Scholarship, and Dr Albert Shimmins Postgraduate Writing-Up Award from the University of Melbourne. I.K.H.L. is supported by the Marsden Fund by the Royal Society of New Zealand (21-UOA-108) and a Project Grant from the Maurice & Phyllis Paykel Trust. I.K.H.L. would like to thank the University of Melbourne for support through the Driving Research Momentum (DRM) initiative. The Walter and Eliza Hall Institute of Medical Research receives funding provided by the NHMRC Independent Research Institute Infrastructure Support Scheme and the Victorian State Government Operational Infrastructure Program.

Monash Proteomics and Metabolomics Platform (MPMP) utilises infrastructure enabled by Bioplatforms Australia (BPA) and the National Collaborative Research Infrastructure Strategy (NCRIS). Additional support was provided to MPMP by Monash eResearch capabilities, including Research Data Storage and Nectar Research Cloud.

## Extended Data Figures

**Extended Data Figure 1.**
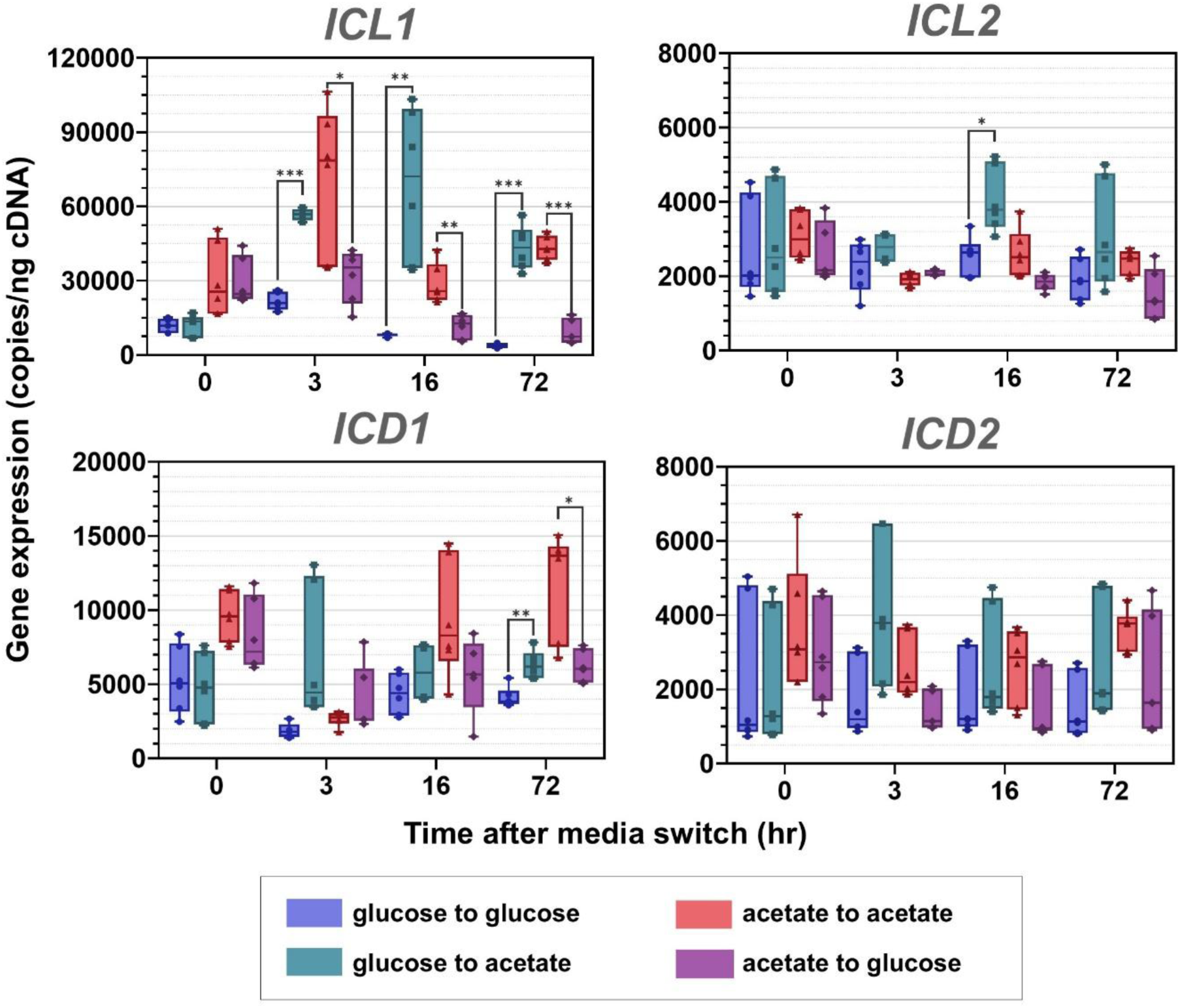
Absolute gene expression of *M. tuberculosis* ICLs and ICDs under different culture conditions. Each data point represents the copy number (normalized to SigA expression) for one replicate. Error bars represent s.d. from six replicates. Asterisks (*) represent a statistically significant difference between the switched and same nutrient control at each time-point using the student’s t-test corrected for multiple comparisons using the Holm-Sidak method: p < 0.001(***), p < 0.01(**), p < 0.05(*).

**Extended Data Figure 2.**
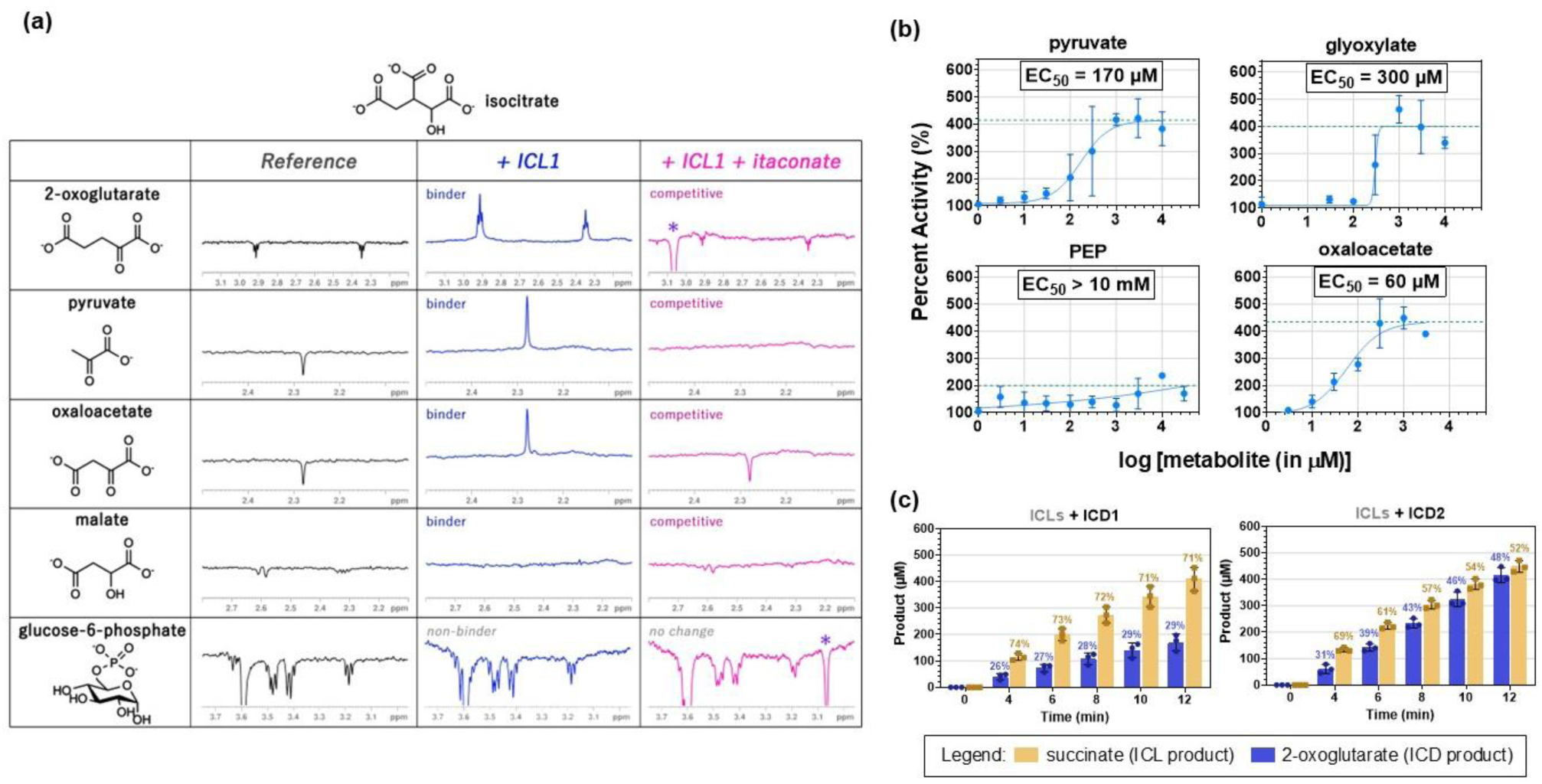
Effect of metabolite ICL inhibitors and ICD2 activators. **(a)** WaterLOGSY spectra of the ICL1 inhibitors in the presence and absence of ICL1 and the active site inhibitor itaconate. A positive or less negative signal relative to the reference spectrum indicates binding. Glucose-6-phosphate (not an ICL inhibitor) was included as a negative control to illustrate the absence of binding. Purple asterisks (*) indicate itaconate peaks. **(b)** Dose response curves of the ICD2 activators. The green dotted line indicates the maximal activation. Error bars represent s.d. from three replicates. **(c)** The relative proportion of ICL (yellow) and ICD (blue) products in the presence of ICLs and only ICD1 or only ICD2. The reaction mixtures contained 42.5 nM of each of the ICLs and 15 nM of ICD1 or ICD2, 5 mM _DL_-isocitrate, 5 mM NADP^+^, 5 mM MgCl_2_, 0.02% NaN_3_, 50 mM Tris-d_11_ in 90% H2O and 10% D2O. Error bars represent s.d. from three replicates.

**Extended Data Figure 3.**
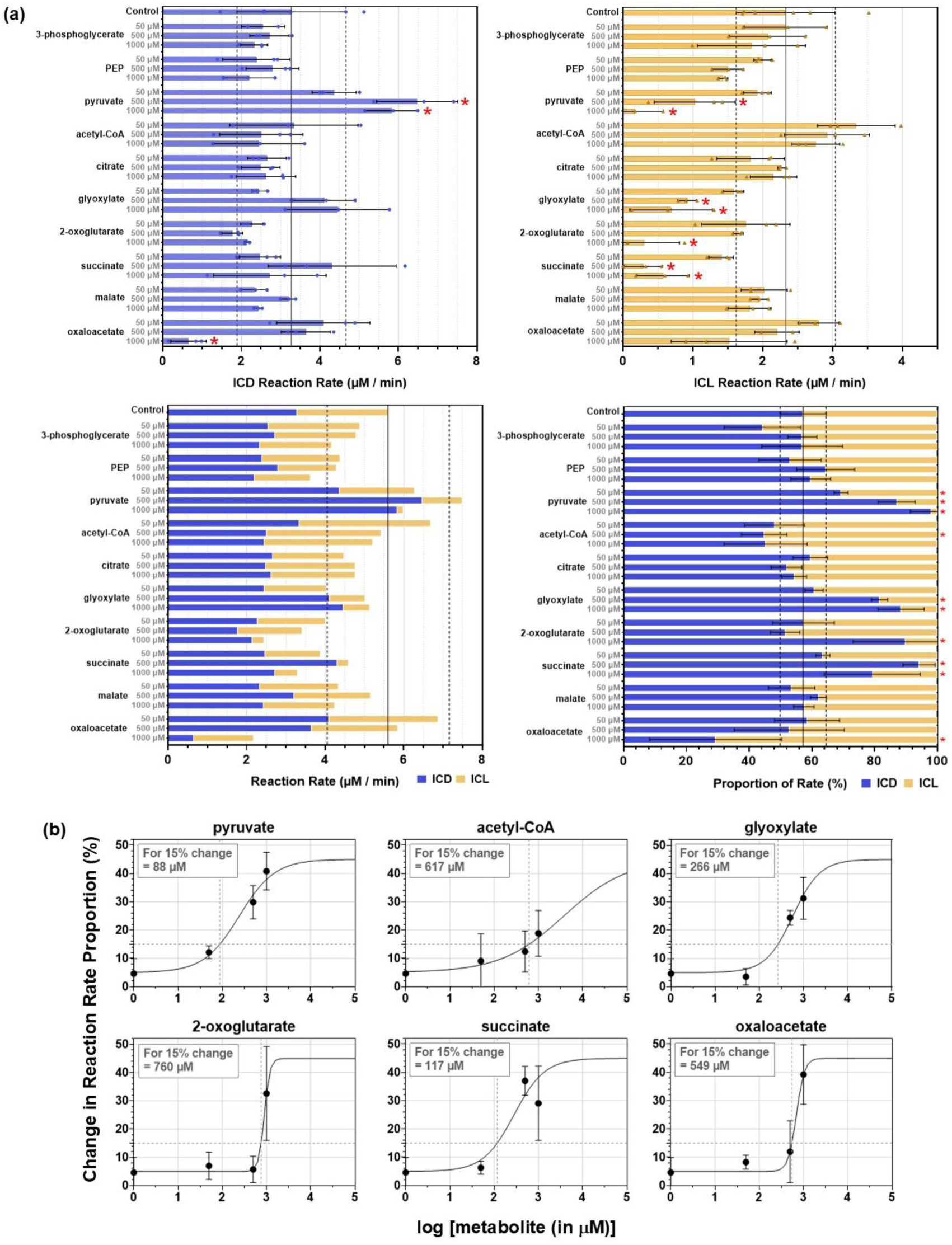
The proportion of the ICD and ICL reaction rate within the total reaction rate under different metabolite effector concentrations. The ICD and ICL reaction rates conditions are given in **(a)**. Error bars indicate s.d. from three replicates. The blue line indicates the value for the control and the error region (for the bottom two graphs). Metabolite conditions that gave a statistically significant difference (p < 0.05) to the control using the student’s t-test were highlighted with a red asterisk. **(b)** Approximate EC_50_ curves for metabolites that gave statistically significant differences in (a). The proportion of flux going through ICD for each data point was subtracted from that of the control, then the absolute value was taken. The proportion of ICD flux in the control was around 55%, so the maximum percent change in flux was set to 45%, and the minimum was set to 5% to represent the error in the control. The dotted line indicates the concentration required to alter the proportion of the rates by 15%.

