## Supplementary Information for "A mixer tap mechanism in *Mycobacterium tuberculosis* regulates a critical metabolic node in response to nutrient availability"

### Supplementary Equations

This equation was taken from Brandt *et al.*[1].

$$\frac{\boldsymbol{v_i}}{\boldsymbol{v_0}} = \frac{\boldsymbol{K_M} + [\boldsymbol{S}]}{\boldsymbol{K_M}\left(\boldsymbol{1} + \frac{[\boldsymbol{I}]}{\boldsymbol{K_i}}\right) + [\boldsymbol{S}]} \quad (\boldsymbol{1})$$

| | | | |
|---|---|---|---|
| $\boldsymbol{v_o}$ | reaction velocity (no inhibitor) | $[\boldsymbol{I}]$ | concentration of inhibitor |
| $\boldsymbol{v_{max}}$ | ligand | $K_M$ | Michaelis constant (substrate concentration at half $v_{max}$) |
| $\boldsymbol{v_i}$ | reaction velocity in the presence of the inhibitor | $K_i$ | inhibition constant |
| $[\boldsymbol{S}]$ | concentration of substrate | | |

At 50% inhibition, $\frac{v_i}{v_0} = 0.5$.

$$K_i = \frac{IC_{50}}{\left(1 + \frac{[S]}{K_M}\right)} \quad (2)$$

For the metabolite screening assay in this study, we want to know the amount of substrate needed to see 50% inhibition with an inhibitor with a $K_i$ of 1 mM if 5 mM inhibitor was used. This means that

$$K_i = 1000\ \mu M$$

$$IC_{50} = 5000\ \mu M$$

### Supplementary Figures

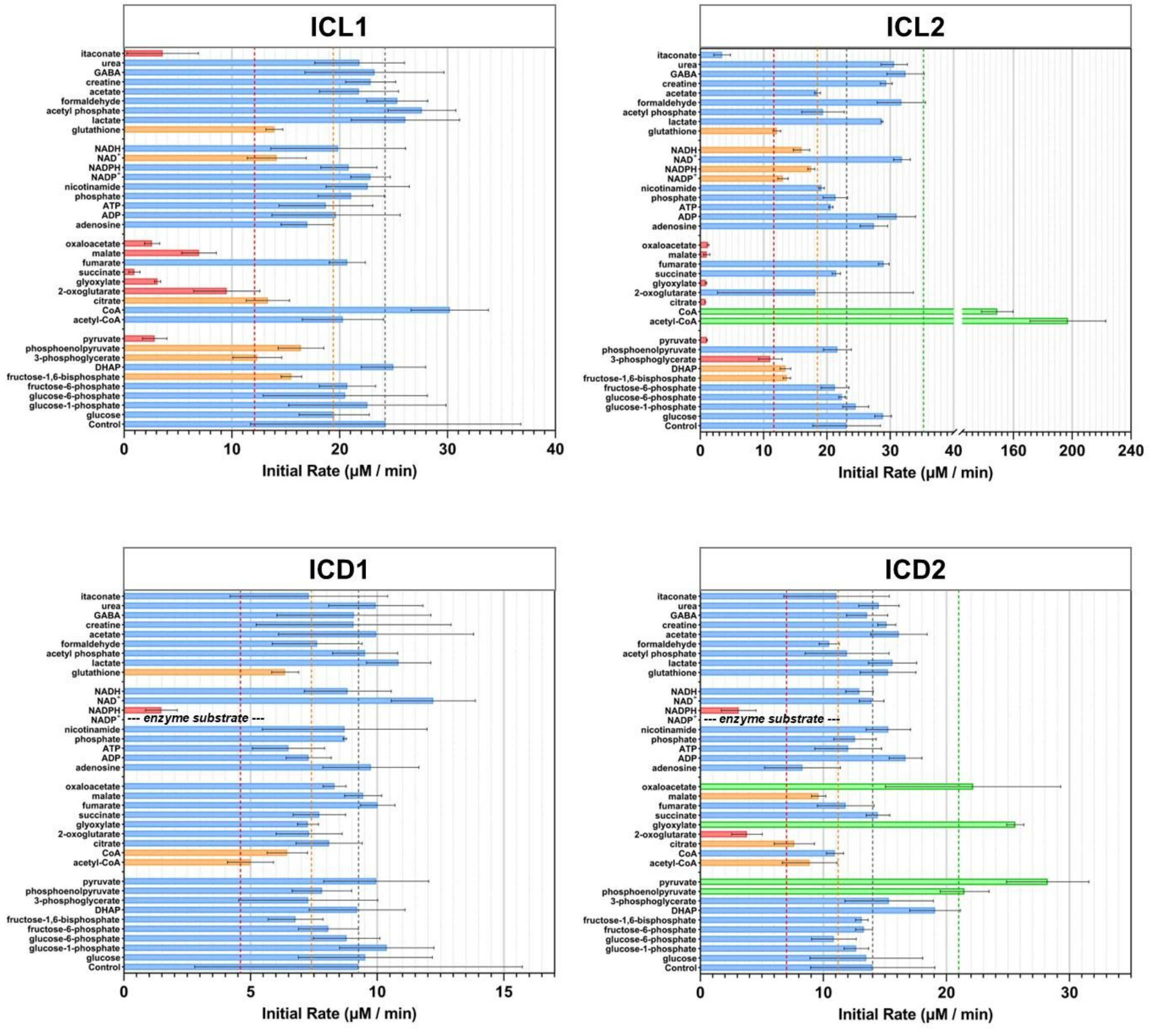

**Supplementary Figure 1. Single concentration metabolite effector screening for *M. tuberculosis* ICLs and ICDs.** The graph shows the initial rate of the ICL/ICD reaction in the presence of different metabolites. Initial rates were measured at <20% substrate turnover. Error bars represent s.d. from three replicates. The metabolites are grouped into four (bottom to top): glycolysis, TCA cycle / glyoxylate shunt, cofactors, others. The reaction mixtures were measured at 25 °C. The reaction mixtures are given in the Supplementary Information. Dotted lines indicated 150% activity (green), 100% activity (gray), 80% activity (orange), and 50% activity (red). Metabolites that gave an activity below 50% (including the error bar) were coloured in red. Metabolites that gave an activity between 50-80% (including the error bar) were coloured in orange, and metabolites that increased the activity beyond 150% was coloured in green.

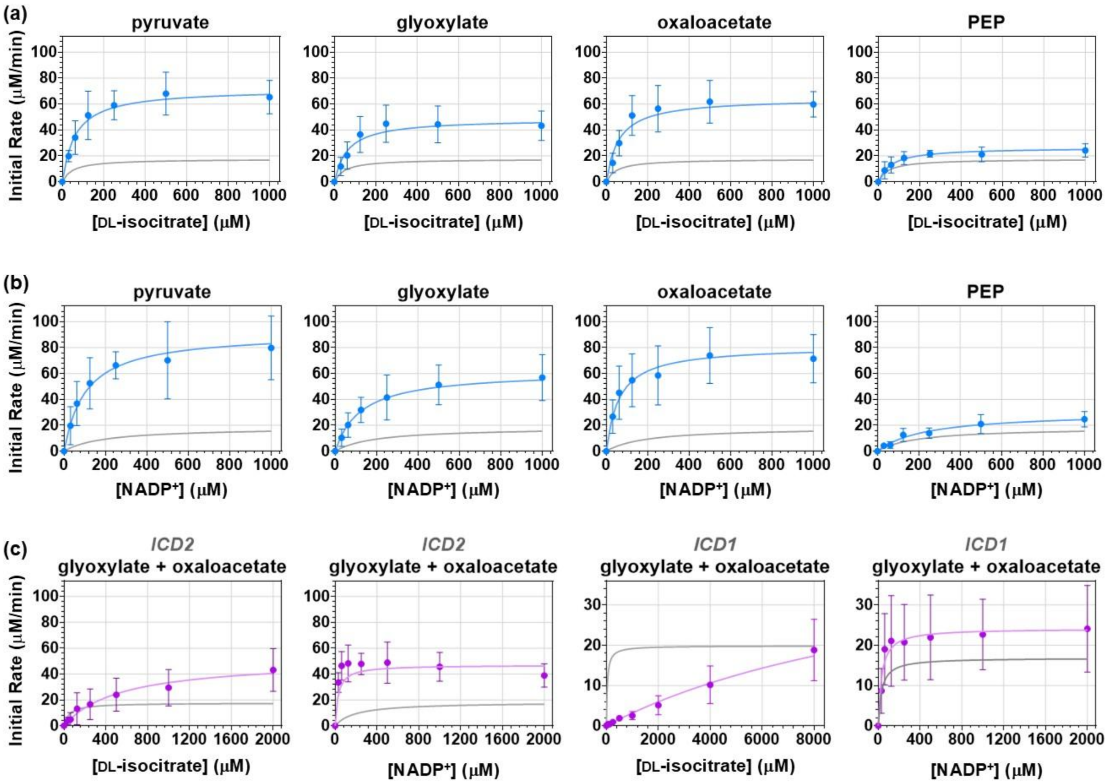

(d)

| *DL-isocitrate* | ICD2 | | | | | | ICD1 | |
|---|---|---|---|---|---|---|---|---|
| | Control | pyruvate | glyoxylate | OAA | PEP | glyoxylate + OAA | Control | glyoxylate + OAA |
| $K_M$ (μM) | 50 ± 36 | 63 ± 26 | 64 ± 51 | 60 ± 52 | 63 ± 31 | 544 ± 388 | 36 ± 10 | 14000 ± 6000 |
| $V_{max}$ (μM/min) | 18 ± 2 | 72 ± 6 | 49 ± 7 | 65 ± 10 | 27 ± 3 | 52 ± 17 | 20 ± 1 | 48 ± 12* |
| [enzyme] | 15 nM | 15 nM | 15 nM | 15 nM | 15 nM | 15 nM | 25 nM | 25 nM |
| $k_{cat}$ ($s^{-1}$) | 20 ± 3 | 80 ± 7 | 54 ± 8 | 72 ± 12 | 29 ± 3 | 57 ± 19 | 13 ± 1 | 32 ± 8 |

(e)

| *NADP+* | ICD2 | | | | | | ICD1 | |
|---|---|---|---|---|---|---|---|---|
| | Control | pyruvate | glyoxylate | OAA | PEP | glyoxylate + OAA | Control | glyoxylate + OAA |
| $K_M$ (μM) | 191 ± 79 | 104 ± 43 | 124 ± 45 | 62 ± 28 | 236 ± 79 | 30 ± 23 | 31 ± 18 | 32 ± 27 |
| $V_{max}$ (μM/min) | 18 ± 1 | 92 ± 7 | 62 ± 4 | 81 ± 6 | 30 ± 2 | 47 ± 6 | 17 ± 2 | 24 ± 3 |
| [enzyme] | 15 nM | 15 nM | 15 nM | 15 nM | 15 nM | 15 nM | 25 nM | 25 nM |
| $k_{cat}$ ($s^{-1}$) | 20 ± 2 | 102 ± 7 | 69 ± 5 | 90 ± 6 | 34 ± 2 | 52 ± 7 | 11 ± 1 | 16 ± 2 |

**Supplementary Figure 2. Michaelis-Menten parameters of *M. tuberculosis* ICDs in the presence of ICD2 activators.** Error bars represent s.d. from three experiments. The Michaelis-Menten curve without the activator is given in gray in all the graphs for comparison as control. Michaelis-Menten curves for ICD2 in the presence of the ICD2 activators are given in **(a)** for DL-isocitrate and **(b)** for NADP+. **(c)** Michaelis-Menten curves of ICD1 and ICD2 in the presence of a mixture of glyoxylate and oxaloacetate. The corresponding Michaelis-Menten parameters are given in **(d)** and **(e)**.

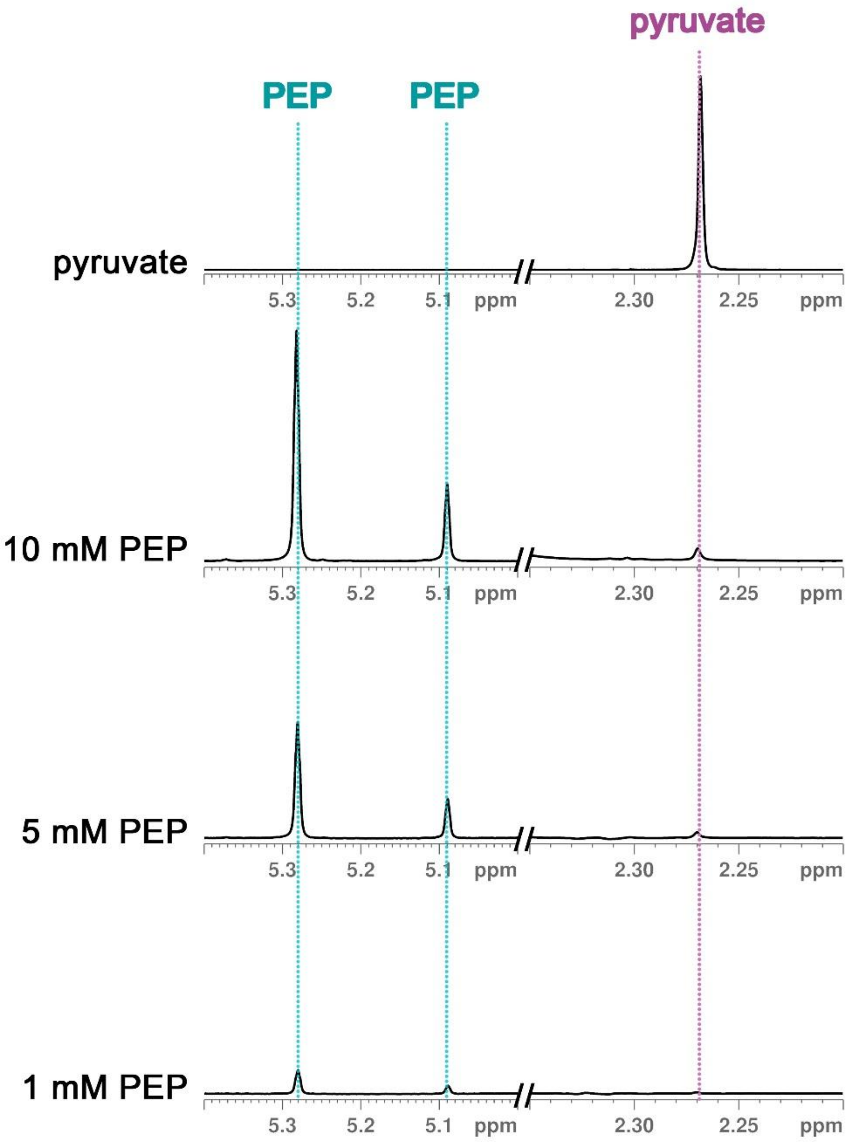

**Supplementary Figure 3. Potential pyruvate impurity present in phosphoenolpyruvate (PEP) samples.** A peak corresponding to the chemical shift of pyruvate was present in PEP samples in a dose-dependent manner. The PEP and pyruvate peaks are labelled in each spectrum. The top spectrum contained 5 mM pyruvate, 50 mM Tris-$d_{11}$, 0.02% $NaN_3$ in 90% $H_2O$ / 10% $D_2O$. The rest of the spectra contained the specified amount of PEP in 50 mM Tris-$d_{11}$, 0.02% $NaN_3$, 90% $H_2O$ / 10% $D_2O$. The pyruvate-only spectrum is not to scale with the rest of the spectra and is only used to illustrate the chemical shift of pyruvate.

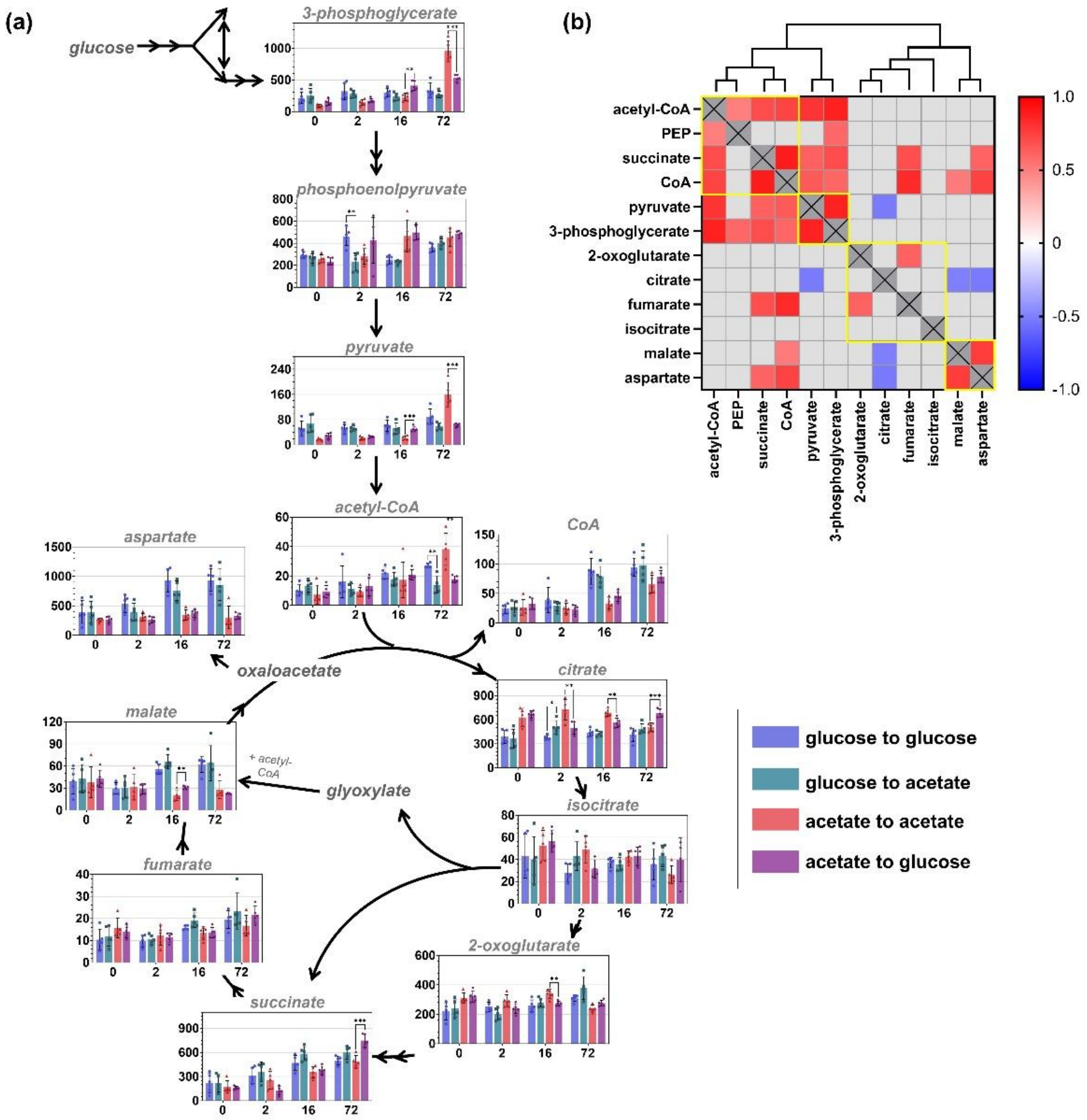

**Supplementary Figure 4. Comparison of metabolite concentrations between *M. tuberculosis* CDC1551 under different culture conditions. (a)** The metabolites were mapped onto the glycolysis and TCA cycle pathways. The x-axis denotes the time point (in hr) after the switch, and the y-axis represents the approximate intracellular concentration of the metabolite (in µM). Error bars represent s.d. from five replicates. Asterisks (*) represent a statistically significant difference to the control using the student's t-test corrected for multiple comparisons using the Holm-Sidak method: $p < 0.001$(***), $p < 0.01$(**), $p < 0.05$(*). Oxaloacetate and glyoxylate were below the limit of detection in all samples, but aspartate was used as a reporter for oxaloacetate. **(b)** Correlation matrix and hierarchical clustering of the glycolytic and TCA cycle metabolites over time under different conditions in the targeted metabolomics experiment. The metabolite concentrations were $\log_{10}$ transformed then Pareto scaled. Correlations that did not show statistical significance ($p > 0.05$) were shaded in grey. The four distinct clusters according to the hierarchical clustering are enclosed in yellow squares.

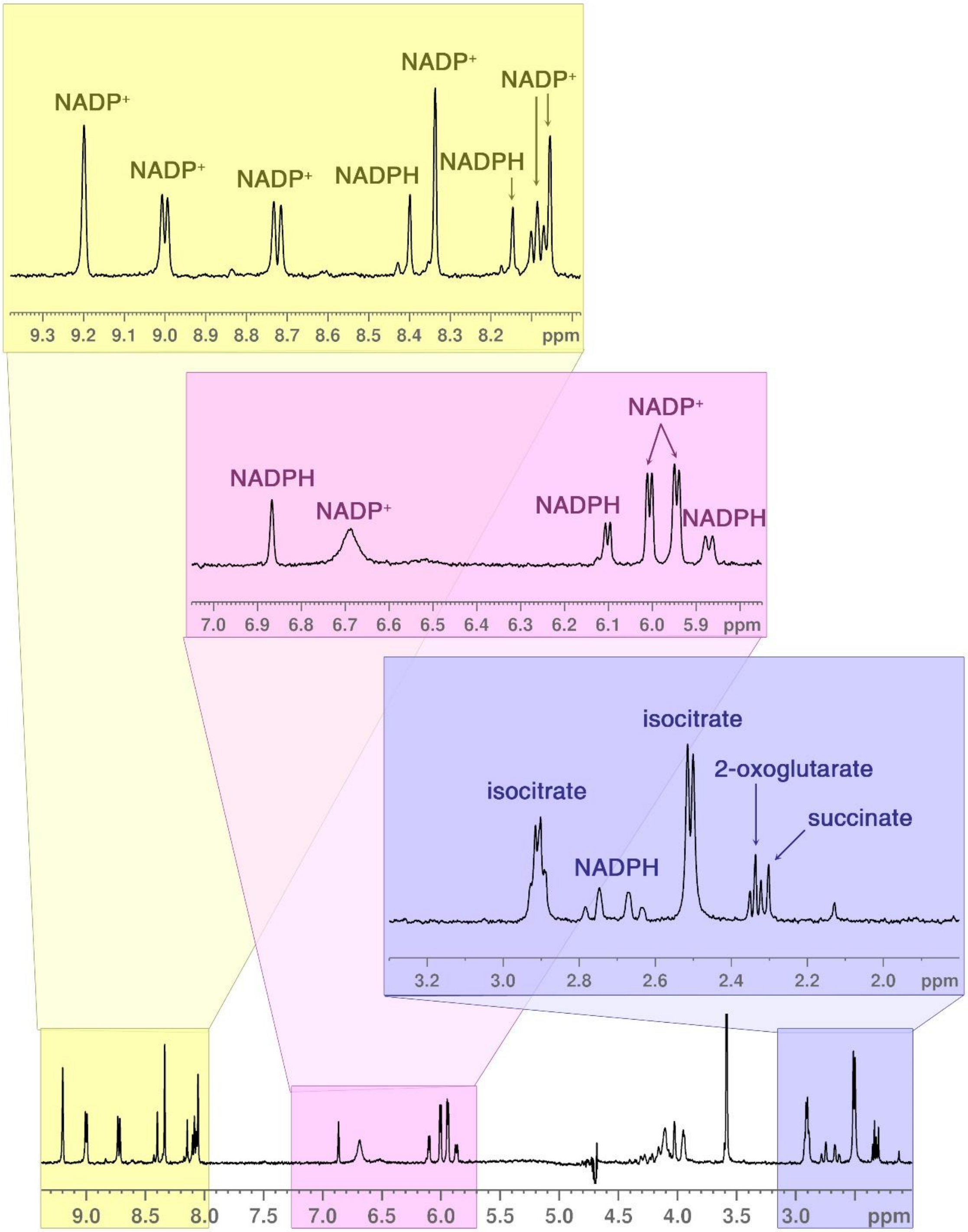

**Supplementary Figure 5. $^1$H NMR spectrum of isocitrate and $NADP^+$ in the presence of both ICL and ICD.** The full spectrum is shown at the bottom, and the different regions of the spectrum is zoomed in to show the substrate and product peaks. The reaction was measured at 25 °C, and the spectrum shown was measured 3.5 min after enzyme addition. The reaction mixture contained 37.5 nM of each of the ICLs and the ICDs, 1 mM DL-isocitrate, 1 mM $NADP^+$, 5 mM $MgCl_2$, 0.02% $NaN_3$, 50 mM Tris-$d_{11}$ in 90% $H_2O$ and 10% $D_2O$.

### Supplementary Tables

**Supplementary Table 1. Calculated and actual substrate $S$ concentrations used for the metabolite screening of *M. tuberculosis* ICLs and ICDs.** A concentration less than the calculated substrate concentration $[S]_{calc}$ is needed to achieve at least 50% inhibition when 5 mM is used for an inhibitor with an inhibition constant of 1 mM.

| | $K_M$ | $[P]_T$ | $[S]_{calc}$ | $[S]_{actual}$ |
|---|---|---|---|---|
| **ICL1** (DL-isocitrate) | 104 μM | 0.2 μM | 416 μM | 250 μM |
| **ICL2** (DL-isocitrate) | 570 μM | 2 μM | 2.3 mM | 1.2 mM |
| **ICD1** (DL-isocitrate) | 143 μM | 0.02 μM | 572 μM | 250 μM |
| **ICD2** (DL-isocitrate) | 103 μM | 0.025 μM | 412 μM | 250 μM |
| **ICD1** ($NADP^+$) | 59 μM | 0.02 μM | 236 μM | 100 μM |
| **ICD2** ($NADP^+$) | 564 μM | 0.025 μM | 2.3 mM | 1 mM |

**Supplementary Table 2. Summary of Michaelis-Menten parameters of *M. tuberculosis* ICLs and ICDs.** Errors indicate s.d. from three separate experiments. The Michaelis-Menten equation was fitted to the data points using Graphpad Prism 8. All measurements were obtained using the absorbance-based assay[2-3]. A range of DL-isocitrate (0 – 2000 μM) and $NADP^+$ (0 – 2000 μM; for ICD) concentrations were tested. The reaction was measured at 25 °C. For testing the parameters for isocitrate, the mixtures contained varying concentrations of DL-isocitrate and 4 mM $NADP^+$ (for the ICDs). For testing the parameters for $NADP^+$ for the ICDs, the mixtures contained varying concentrations of $NADP^+$ and 4 mM DL-isocitrate. All reaction mixtures also had the specified amount of enzyme, 5 mM $MgCl_2$, and 10 mM phenylhydrazine (for ICLs) in 50 mM Tris (pH 7.5).

| | ICL1 | ICL2 | ICD1 | ICD2 |
|---|---|---|---|---|
| ***DL-isocitrate*** | | | | |
| $K_M$ (μM) | 104 ± 19 | 570 ± 130 | 143 ± 31 | 103 ± 45 |
| $V_{max}$ (μM/min) | 22 ± 1 | 20 ± 3 | 31 ± 2 | 44 ± 4 |
| [enzyme] | 200 nM | 2 μM | 25 nM | 50 nM |
| $k_{cat}$ ($s^{-1}$) | 1.8 ± 0.1 | 0.2 ± 0.1 | 20.9 ± 1.2 | 14.7 ± 1.4 |
| ***$NADP^+$*** | | | | |
| $K_M$ (μM) | -- | -- | 59 ± 14 | 564 ± 185 |
| $V_{max}$ (μM/min) | -- | -- | 39 ± 2 | 55 ± 5 |
| [enzyme] | -- | -- | 25 nM | 50 nM |
| $k_{cat}$ ($s^{-1}$) | -- | -- | 26.3 ± 1.3 | 18.2 ± 1.5 |

**Supplementary Table 3. Enzyme concentrations used to test the effect of gene expression on the ICL-ICD flux.** The ratios between the isoforms were inferred from the gene expression data in Extended Data Fig. 1. Enzyme concentrations were given in nM, and the concentrations were scaled according to the ratios so that the lowest concentration of an isoform was 1.5 nM ([ICD2] in condition G72).

| | **Enzyme Concentrations (in nM)** | | | | | | | | | | | | | | | |
|---|---|---|---|---|---|---|---|---|---|---|---|---|---|---|---|---|
| | **glucose to glucose** | | | | **glucose to acetate** | | | | **acetate to acetate** | | | | **acetate to glucose** | | | |
| | **0 h** | **2 h** | **16 h** | **72 h** | **0 h** | **2 h** | **16 h** | **72 h** | **0 h** | **2 h** | **16 h** | **72 h** | **0 h** | **2 h** | **16 h** | **72 h** |
| ICL1 | 11.6 | 21.2 | 8.0 | 3.9 | 11.9 | 55.8 | 68.1 | 42.8 | 29.8 | 70.2 | 28.3 | 42.5 | 29.1 | 31.4 | 11.4 | 9.3 |
| ICL2 | 2.6 | 2.2 | 2.5 | 1.9 | 2.9 | 2.7 | 4 | 3 | 3.0 | 1.9 | 2.6 | 2.3 | 2.6 | 2.1 | 1.8 | 1.5 |
| ICD1 | 5.2 | 1.8 | 4.3 | 4.1 | 4.7 | 6.7 | 5.7 | 6.2 | 9.4 | 2.6 | 9.3 | 11.7 | 8.1 | 3.9 | 5.5 | 6.1 |
| ICD2 | 2.2 | 1.7 | 1.8 | 1.5 | 2.1 | 4 | 2.6 | 2.7 | 3.6 | 2.6 | 2.6 | 3.6 | 2.9 | 1.4 | 1.5 | 2.3 |
| Total | 21.6 | 27.0 | 16.5 | 11.4 | 21.6 | 69.2 | 80.4 | 54.7 | 45.9 | 77.3 | 42.8 | 60.1 | 42.7 | 38.8 | 20.2 | 19.2 |
| Percent ICLs | 66% | 87% | 63% | 51% | 69% | 85% | 90% | 84% | 72% | 93% | 72% | 75% | 74% | 86% | 65% | 56% |
| Percent ICDs | 34% | 13% | 37% | 49% | 31% | 15% | 10% | 16% | 28% | 7% | 28% | 25% | 26% | 14% | 35% | 44% |

**Supplementary Table 4. Metabolite cocktails used to test the effect of metabolite mixtures on the ICL-ICD flux.** The metabolite cocktails were based on the estimated intracellular concentrations (in µM) measured in the metabolomics study in Supplementary Fig. 4a. Abbreviations: 3-PG = 3-phosphoglycerate; 2-OG = 2-oxoglutarate; OAA = oxaloacetate.

| | **Metabolite Concentrations (in µM)** | | | | | | | | | | | | | | | |
|---|---|---|---|---|---|---|---|---|---|---|---|---|---|---|---|---|
| | **glucose to glucose** | | | | **glucose to acetate** | | | | **acetate to acetate** | | | | **acetate to glucose** | | | |
| | **0 h** | **2 h** | **16 h** | **72 h** | **0 h** | **2 h** | **16 h** | **72 h** | **0 h** | **2 h** | **16 h** | **72 h** | **0 h** | **2 h** | **16 h** | **72 h** |
| 3-PG | 200 | 300 | 300 | 300 | 250 | 300 | 250 | 250 | 100 | 150 | 250 | 1000 | 150 | 150 | 400 | 550 |
| PEP | 300 | 450 | 250 | 350 | 250 | 250 | 250 | 400 | 250 | 300 | 450 | 450 | 250 | 450 | 500 | 500 |
| pyruvate | 50 | 50 | 60 | 90 | 70 | 50 | 50 | 60 | 20 | 20 | 20 | 160 | 30 | 25 | 50 | 60 |
| acetyl-CoA | 10 | 15 | 20 | 30 | 15 | 10 | 20 | 15 | 5 | 10 | 20 | 40 | 10 | 15 | 20 | 20 |
| CoA | 25 | 40 | 90 | 100 | 30 | 30 | 80 | 100 | 25 | 20 | 30 | 70 | 30 | 20 | 45 | 80 |
| citrate | 400 | 400 | 450 | 400 | 350 | 500 | 400 | 500 | 650 | 700 | 700 | 500 | 650 | 500 | 550 | 700 |
| 2-OG | 200 | 250 | 250 | 300 | 250 | 200 | 300 | 400 | 300 | 300 | 350 | 250 | 300 | 250 | 250 | 250 |
| succinate | 200 | 300 | 450 | 500 | 200 | 350 | 600 | 600 | 150 | 250 | 350 | 500 | 150 | 100 | 400 | 750 |
| fumarate | 10 | 10 | 15 | 20 | 10 | 10 | 20 | 25 | 15 | 10 | 15 | 15 | 15 | 10 | 15 | 20 |
| malate | 40 | 30 | 55 | 60 | 40 | 30 | 65 | 65 | 40 | 30 | 20 | 30 | 40 | 30 | 30 | 20 |
| OAA | 400 | 550 | 950 | 950 | 400 | 400 | 750 | 850 | 250 | 300 | 350 | 300 | 250 | 250 | 350 | 300 |

**Supplementary Table 5. Expression conditions for all proteins used in this study.**

| **Protein** | **Plasmid Backbone** | **Expression System (*E.coli* strain)** | **Media** | **[IPTG]** | **Inductio n Temp.** |
|---|---|---|---|---|---|
| ICL1 | pNIC28-Bsa4 | BL21 (DE3) | 2xYT | 0.2 mM | 18 °C |
| ICL2 | pYUB28b | BL21 LOBSTR (+pGro7) | Terrific Broth (TB) | 0 mM (leaky) | 37 °C |
| ICD1 | pET28a(+) | BL21 LOBSTR (+pGro7) | 2xYT | 0.2 mM | 18 °C |
| ICD2 | pET28a(+) | BL21 (DE3) | LB | 0.2 mM | 18 °C |

**Supplementary Table 6. Buffers used for protein purification.**

| | Buffer A | Buffer B | Storage Buffer |
|---|---|---|---|
| ICL1, ICD1 | 50 mM HEPES<br>500 mM NaCl<br>5 mM imidazole<br>pH 7.8 | 50 mM HEPES<br>500 mM NaCl<br>500 mM imidazole<br>pH 7.8 | 50 mM Tris<br>150 mM NaCl<br>pH 7.5 |
| ICD2 | 50 mM HEPES<br>150 mM NaCl<br>5 mM imidazole<br>pH 7.8 | 50 mM HEPES<br>150 mM NaCl<br>500 mM imidazole<br>pH 7.8 | 50 mM Tris<br>150 mM NaCl<br>pH 7.5 |
| ICL2 | 50 mM HEPES<br>150 mM NaCl<br>5 mM imidazole<br>1 mM β-mercaptoethanol<br>pH 7.5 | 50 mM HEPES<br>150 mM NaCl<br>500 mM imidazole<br>1 mM β-mercaptoethanol<br>pH 7.5 | 50 mM HEPES<br>150 mM NaCl<br>pH 7.5 |
